# Choosing the best or avoiding the worst: complementary and opponent value signals in the human brain

**DOI:** 10.1101/2025.11.04.686111

**Authors:** Clarissa Baratin, Alizée Lopez-Persem, Philippe Kahane, Lorella Minotti, Jiri Hammer, Petr Marusic, Anca Nica, Sylvain Rheims, Louis Maillard, Marie Denuelle, Emmanuel Barbeau, Blandine Chanteloup-Foret, Mathias Pessiglione, Guillaume Becq, Julien Bastin

## Abstract

Whether the brain encodes the subjective value of pleasant and unpleasant situations via a single integrated circuit or through distinct pathways remains unresolved. It is also unclear how neural value signals guide choices when the context involves choosing the most favorable option or avoiding the most unfavorable one. To address these questions, we recorded intracerebral activity from 27 participants as they first rated pleasant and unpleasant scenarios and then made binary choices across all combinations of item domain (pleasant/unpleasant) and choice frame (choose best/worst). Ventromedial prefrontal cortex (vmPFC) broadband gamma activity (BGA, 50–150 Hz) tracked the value of pleasant items during rating and pre-choice, correlating positively with the most pleasant item irrespective of choice frame. Similarly, anterior insula (aIns) BGA tracked the value of unpleasant items during rating and pre-choice, correlating negatively with the most unpleasant item irrespective of choice frame. The difference in relative value correlations between the vmPFC and aIns was larger for choices involving a conflict between item domain and choice frame, consistent with slower response times and choices less aligned with value differences in these trials. These findings reveal dissociable yet complementary neural systems for pleasant and unpleasant valuation and highlight how such signals may guide flexible decision-making across contexts.

## Introduction

How does the human brain adapt value computations to guide choice across appetitive and aversive contexts? A central question in decision neuroscience is whether the brain computes value using a single, valence-general system or relies on parallel appetitive and aversive circuits. Adding to this uncertainty, the properties of value signals may vary with task context, for instance when evaluating options in isolation versus during choice.

Extensive human neuroimaging work (with functional magnetic resonance imaging, fMRI) has identified reliable value-related activity in the ventromedial prefrontal cortex (vmPFC), ventral striatum, and posterior cingulate cortex (Kable and Glimcher 2007; Lebreton et al. 2009; Liu et al. 2011; Bartra et al. 2013; Clithero and Rangel 2014). Within this “valuation system,” vmPFC responses scale positively with the value of appetitive options and negatively with aversive ones, consistent with a common currency model (Tom et al. 2007; Plassmann et al. 2010; Bartra et al. 2013; Lindquist et al. 2016). However, parallel evidence suggests that aversive valuation may instead rely on distinct regions such as the anterior insula (aIns) or dorsomedial prefrontal cortex (Corradi-Dell’Acqua et al. 2016; Le Bouc and Pessiglione 2022), with opposing activity patterns in the ventral striatum and aIns during risk-taking (Kuhnen and Knutson 2005). In monkeys, pioneering single-cell studies reported that lateral orbitofrontal cortex (lOFC) neurons integrate both appetitive and aversive values (Hosokawa et al. 2007; Morrison and Salzman 2009), but direct comparisons between stimulus domains are confounded by sensory differences between rewards (juice) and punishments (air puff or shock). Token-based paradigms mitigate this confound, revealing that the aIns contains more neurons encoding losses than gains (Yang et al. 2022), whereas orbitofrontal (OFC) neurons encode value across both outcomes (Tang et al. 2024), leaving unresolved whether appetitive and aversive values are represented within a single system or distinct circuits.

Beyond anatomical uncertainties, human and animal studies provide contrasting evidence regarding the sign of value coding. In humans, vmPFC activity is consistently positively correlated with value in fMRI studies (Bartra et al. 2013). In monkeys, both single neurons and broadband gamma activity (BGA, 50–150 Hz) show heterogeneous coding, with either positive or negative correlations with value in the OFC (Rich and Wallis 2017; Tang et al. 2024; Sharma et al. 2025). Recently, studies have sought to bridge the gap between human functional imaging and monkey single-cell experiments by examining value signals in the BGA of epileptic patients who had intracerebral electrodes implanted. This approach is particularly valuable because BGA provides a time-resolved index of local neuronal activity that correlates positively with both hemodynamic responses and local spiking activity (Mukamel et al. 2005; Niessing 2005; Lachaux et al. 2007; Nir et al. 2007; Manning et al. 2009). Importantly, intracerebral recordings in humans show that, when statistically assessed across all recording sites within a region, BGA–value correlations are predominantly positive throughout both the lateral and ventromedial OFC (Lopez-Persem et al. 2020; Shih et al. 2023), providing a direct human neural measure that can link population-level fMRI signals with the cellular-level coding observed in monkeys. Moreover, these studies focused exclusively on appetitive events, leaving it unclear how BGA encodes the value of aversive events. This question is important, since prior intracerebral evidence suggests opponent BGA responses in the vmPFC and aIns for appetitive and aversive stimuli, respectively (Gueguen et al. 2021; Cecchi et al. 2022). Therefore, the first aim of this study was to determine how human BGA represents the value of single options across appetitive and aversive contexts. To this end, intracerebral recordings were acquired while epileptic patients rated the (un)pleasantness of single text-based scenarios describing pleasant or unpleasant life events.

The second aim of this study was to determine how value comparison occurs during binary decisions and whether it depends on item domain (appetitive vs. aversive) and choice frame (selecting the best or worst option). vmPFC activity is often interpreted as encoding relative value during binary choices in humans, positively for chosen and negatively for unchosen options (Hunt et al. 2012; Nicolle et al. 2012; Boorman et al. 2013). Yet, alternative evidence suggests a sum-based signal (Clairis and Pessiglione 2022) or goal-aligned coding depending on choice frame (Frömer et al. 2019; Sepulveda et al. 2020; Castegnetti et al. 2021). Although Pavlovian links between appetitive–approach and aversive–avoidance behaviors are well established (Boureau and Dayan 2011), their neural counterparts remain poorly understood, particularly regarding how item domain interacts with choice frame. We therefore asked patients to perform binary decisions under a factorial design crossing stimulus domain (pleasant vs. unpleasant) and choice frame (“choose best” vs. “choose worst”), while recording intracerebral activity to reveal how the brain encodes value across these contexts. In the pleasant domain, in typical choices, we tend to approach the most pleasant item, but in the “choose worst” condition participants were instructed to select the least pleasant one, thereby dissociating the value of the chosen option from that of the most pleasant item. In contrast, in the unpleasant domain, choices often involve avoiding the most unpleasant item, but in the “choose best” condition participants were instructed to select the least unpleasant one, again dissociating the value of the chosen option from that of the most unpleasant item.

## Results

### Behavior

27 participants performed valuation and choice tasks while under stereo-electroencephalographic (sEEG) monitoring (Fig. 1a; age 33.6 ± 1.9 years old, 16 females, see demographic and clinical details Table S1). During the valuation task, which was performed first, participants were asked to rate pleasant and unpleasant hypothetical situations on a scale of how much they liked (for pleasant items) or disliked (for unpleasant items) each of them. In total, 120 pleasant and 120 unpleasant items were presented one by one in alternating pleasant and unpleasant blocks (each block consisting of 20 item ratings, see Methods). Although during the experiments the scale changed from "I like it…" in pleasant trials to "I dislike it…" in unpleasant trials, in the following, higher ratings always represent more pleasant values, and lower ratings always represent more unpleasant values. Unsurprisingly, during pleasant blocks, a majority of ratings was positive whereas during unpleasant blocks, a majority of ratings was negative (Fig. 1b). We also found that participants took slightly longer to rate unpleasant items than pleasant items (Fig. 1c; Unpleasant response time = 5.39 ± 0.30 s; Pleasant response time = 5.04 ± 0.25 s; t_(26)_ = 3.29, p = 0.0029; paired two-tailed t-test).

**Figure 1.**
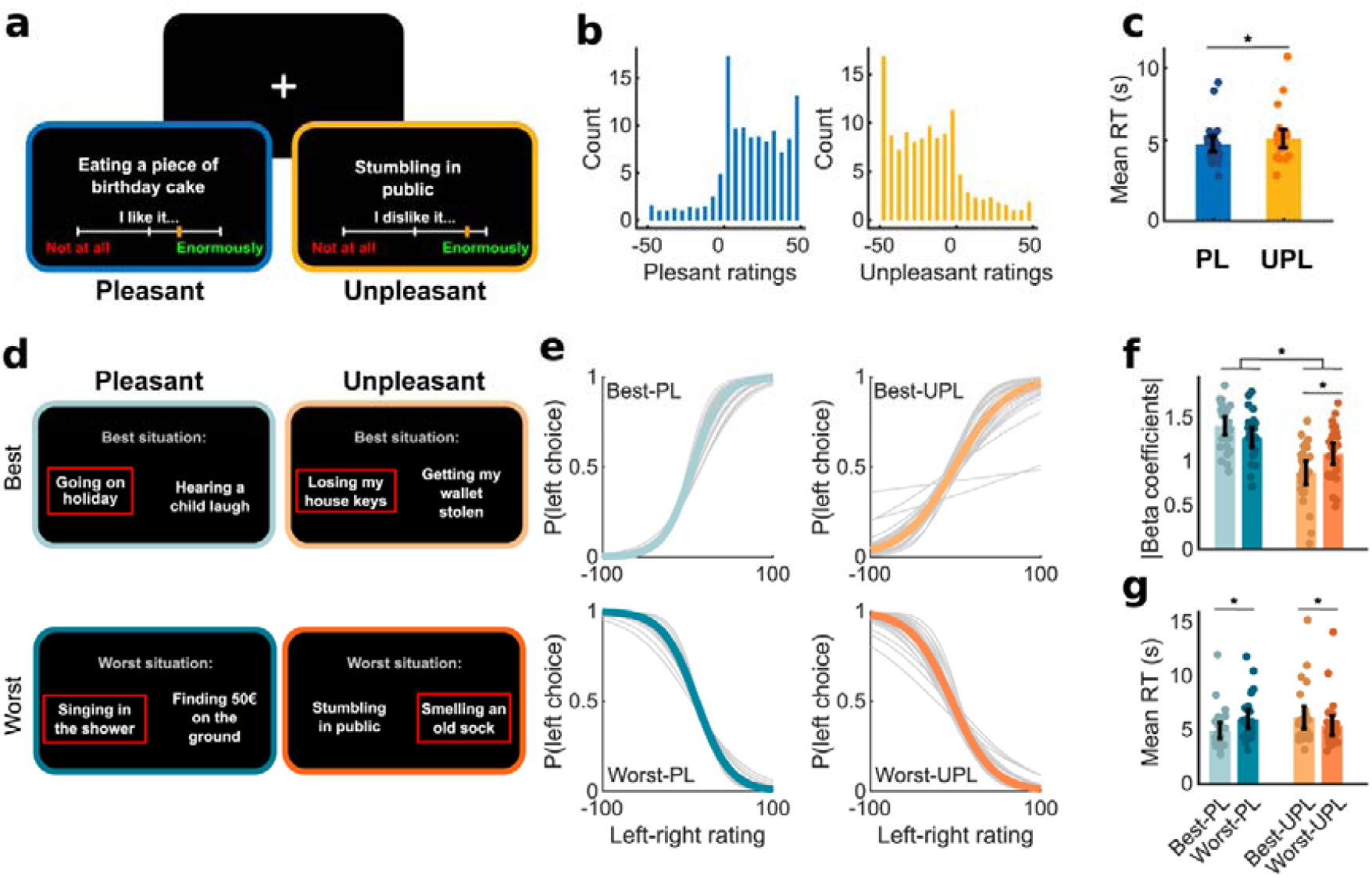
Experimental setup, tasks and behavioral results. (a) During the valuation task, participants rated pleasant and unpleasant situations on a scale of how much they liked (for pleasant items) or disliked (for unpleasant items) each of them. (b) Distributions of value ratings across conditions and participants (c) Mean ± SEM response time across participants for pleasant (PL) and unpleasant (UPL) rating trials. (d) During the four blocks of choices, participants chose between two options, with trials organized in a 2x2 factorial design (2 stimulus domain: pleasant, unpleasant by 2 choice frame: best, worst). (e) Choice behavior. Generalized mixed-effects models were used to predict the probability of choosing the left item as a function of the difference between left and right item ratings. Colored lines represent fixed effects. Grey lines represent random effects (one line per subject). (f) Comparison of random-effect regression coefficients (absolute values) extracted from the choice behavior models represented in (e). (g) Mean response time across participants for the four blocks of choices. In bar plots (f-g), each dot represents a participant, and error bars represent the SEM across participants. Asterisks indicate significance (p < 0.05).

During the binary choice tasks (Fig. 1d) two items were presented on every trial and participants were instructed to choose either their most preferred (“choose best”) or least preferred (“choose worst”) item. Trials were organized and blocked in a 2×2 factorial design (pleasant/unpleasant items by choice goal: best/worst), with each of the four resulting blocks consisting of 60 choice trials. Option pairings were performed based on the ratings obtained from the valuation task, trials were randomized within blocks, and block order was counterbalanced across participants (see Methods). To test whether participants’ choices could be predicted from their value ratings (Fig. 1e), we ran generalized mixed-effects models with participant as a random factor for each choice block separately, using the difference between left and right option ratings to predict a left choice. In the "pleasant - choose best" condition for example, a high rating difference between left and right options (i.e. the left option had a higher hedonic value than the right) should predict a left choice, while in the "unpleasant - choose worst" condition, a high rating difference between left and right options (i.e. the left option had a higher hedonic value than the right) should predict a right choice. We found that rating differences significantly explained choices when participants were asked to choose the best of two pleasant situations (β = 1.41 ± 0.12, t_(1603)_ = 12.11, p = 2.10^−32^), or to choose the worst of two pleasant situations (β = -1.27 ± 0.12, t_(1574)_ = -10.77, p = 4.10^−26^). Similarly, rating differences significantly explained choices when participants were asked to choose the best of two unpleasant situations (β = 0.87 ± 0.11, t_(1581)_ = 7.98, p = 3.10^−15^) or to choose the worst of two unpleasant situations (β = -1.09 ± 0.11, t_(1607)_ = -9.59, p = 3.10^−21^).

We next investigated behavioral differences between choices as a function of item domain (pleasant vs. unpleasant) and choice frame (best vs. worst). First, regression coefficients from the mixed-effects models were used to index choice consistency, reflecting how strongly choices depended on the value difference between items (Fig. 1f). Choice consistency was compared across blocks, using a 2x2 ANOVA with stimulus domain and choice frame as within-participant factors. We found a main effect for stimulus domain (F_(1,26)_ = 65.04, p = 2.10^−8^) and an interaction effect between stimulus domain and choice frame (F_(1,26)_ = 15.09, p = 6.10^−4^). Post-hoc comparisons (Tukey) revealed that choices were more consistent when options were pleasant rather than unpleasant (p < 0.05). Furthermore, in the case of unpleasant options, choices were better predicted by value ratings when participants were asked to choose the worst option rather than the best one (p < 0.05). Second, we compared reaction times (RTs) across contexts (Fig. 1g) and found a main effect for item domain (F_(1,26)_ = 4.32, p = 0.048; 2x2 within-participants ANOVA) and an interaction effect between domain and choice frame (F_(1,26)_ = 51.1, p = 1.10^−7^). Post-hoc comparisons (Tukey) revealed that participants were faster when asked to choose the best option during pleasant choices (p = 2.10^−5^) and faster when asked to choose the worst option during unpleasant choices (p = 0.001). Taken together, choice consistency and RT analyses results indicate that choosing the better of two pleasant items or the worst of two unpleasant items were the most natural conditions for subjects, recalling the existence of Pavlovian links between appetitive items and approach behavior and between aversive items and avoidance behavior (Boureau and Dayan 2011).

Finally, we performed an exploratory analysis to check that RT decreased with overall value when asked to choose the best option and instead increased with overall value when asked to choose the worst, as previously shown (Frömer et al. 2019). Specifically, we used generalized linear mixed-effects models run per block to test how overall value predicted RT and indeed found that the relationship between overall value and RT depended on the task goal (Fig. S1).

### sEEG results

The sEEG dataset comprised 3279 recording sites (i.e., each site corresponding to a bipolar contact-pair) from 26 patients (one fewer than in the behavioral dataset due to a technical issue, see Methods). For each patient, sites were localized on the native anatomical scan and labeled according to the MarsAtlas parcellation scheme (Auzias et al. 2016). After excluding artifact-contaminated sites, 2671 gray-matter sites across 41 parcels (≥ 15 sites per parcels with ≥ 5 patients each) were included in further analyses (Fig. 2).

**Figure 2.**
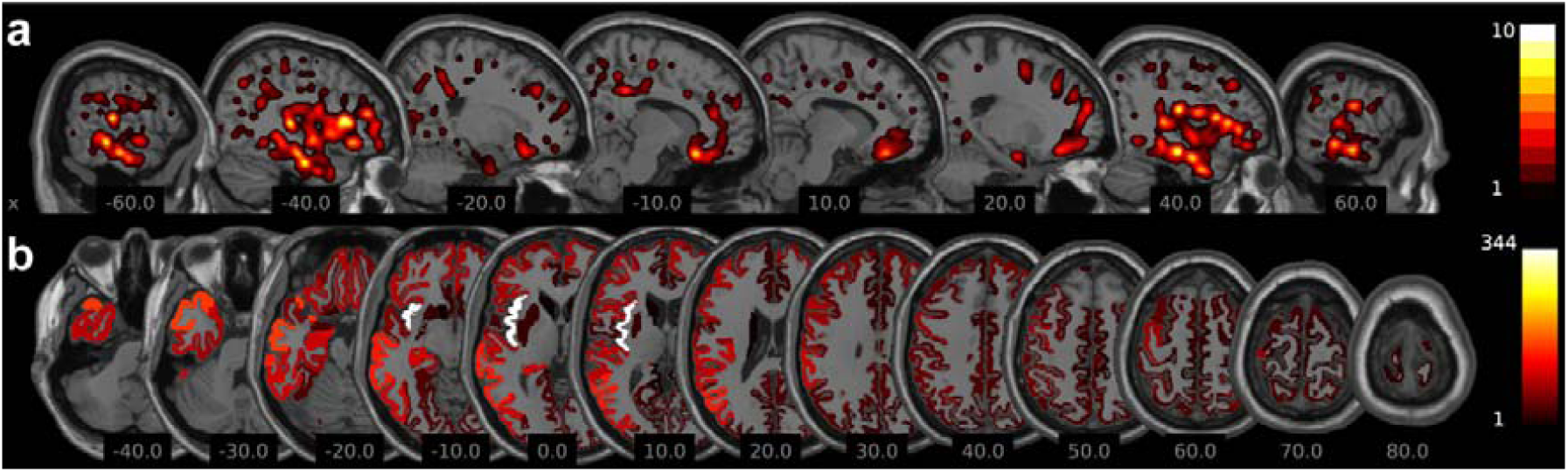
Anatomical locations of recording sites. (a) Sagittal slices of a brain template on which were superimposed the anatomical locations of recording sites across participants (n = 26). Densities are estimated using a Gaussian kernel with a standard deviation of 3 mm for each recording site. (b) Axial slices of a brain template representing all brain parcels (MarsAtlas, see Methods). Color coding (from dark red to white) indicates the number of recording sites in each brain area after pooling left and right sides. The x (a) and z (b) coordinates refer to the MNI (Montreal Neurological Institute) coordinates of the slice displayed.

We initially focused sEEG analyses on BGA as a neural proxy of local neuronal activity, due to its close association with both fMRI and population-level neuronal spiking. This choice was also justified given repeated demonstrations with large datasets that lower frequencies in sEEG signals are redundant when signaling a variety of variables, such as stimulus value, prediction errors or mood fluctuations (Lopez-Persem et al. 2020; Cecchi et al. 2021; Gueguen et al. 2021).

To determine whether a brain region was significantly correlated with the value of pleasant and/or unpleasant items, we first averaged BGA over the interval from stimulus onset to the participant’s response in each trial. This time window was selected because it corresponds to the period during which the brain is expected to evaluate item value. For each site within a region of interest, we then performed a regression of trial-by-trial BGA against item value, yielding one regression estimate per site and stimulus domain. Significance was assessed across all sites within each region by testing the regression estimates against chance level (Student’s t-test, two-sided, corrected for multiple comparisons across regions). Notably, no brain region showed a sustained correlation with both pleasant and unpleasant item values. In contrast, three regions showed a positive correlation with the value of pleasant items: the vmPFC (t[[ = 7.05, p = 6.10^−10^), posterior cingulate cortex (pCC; t[[ = 4.75, p = 3.10^−5^), and caudo-medial visual cortex (cmVC; t[[ = 3.78, p = 6.10^−4^) and two other brain areas showed a negative correlation with the value of unpleasant items: the aIns (t_166_ = -4.11, p = 6.10^−5^) and rostro-medial visual cortex (rmVC: t_32_ = -3.61, p = 3.10^−5^).

To investigate the temporal dynamics of value coding within these ROIs, we performed time-resolved regressions. For each site and each time point, we regressed BGA against item value across trials, for pleasant and unpleasant items separately, producing a time series of regression estimates for each site. To assess significance at the regional level, we tested these estimates across sites using two-sided, one-sample Student’s t-tests. Multiple comparisons across time points were controlled for using a non-parametric cluster-level correction, and a Bonferroni adjustment was applied for the 42 ROIs tested (see Methods). Analyses were time-locked to both stimulus onset (−0.5 to 5 s) and response onset (−5 to 0.5 s) to cover the full valuation period. Since the posterior cingulate and caudo-medial visual cortices — for pleasant items (Fig. S2) — and rostro-medial visual cortex — for unpleasant items (Fig. S3) — showed later and weaker value-related signals compared to the two other brain regions (vmPFC and aIns), we focused subsequent analyses on the latter to characterize value coding.

In the vmPFC, we found that most sites showed a significant positive correlation between average BGA (measured between stimulus and response onset) and pleasant value (Fig. 3a; n = 80 vmPFC sites from 13 subjects). In a typical vmPFC site, BGA responses increased with item value among pleasant items, with significantly stronger activity for highly rated pleasant items compared to those rated as less pleasant (Fig. 3b; p < 0.05). Averaging BGA from stimulus onset to response, this vmPFC site showed a significant correlation with pleasant p < 0.05), but not unpleasant, value (Fig. 3c). This electrophysiological pattern was consistently observed, with vmPFC BGA correlating positively with the value of pleasant items across all sites (1.1–2.1 s, sum(t[[) = 423.2, p_corr_ < α_c_; Fig. 3d). The correlation was stronger for pleasant than unpleasant items (Fig. 3d; 1.4–2 s, sum(t[[) = 230.7, p_corr_ < α_c_), and similar effects were observable prior to response onset (Fig. 3d). Critically, the oisitive correlaton between BGA and pleasant value observed in the vmPFC was not present in neighboring orbitofrontal subregions. We found either no significant correlation with pleasant value in the ventro-medial OFC (n = 51 sites) and ventral OFC (n = 82 sites) or a negative correlation in the ventro-lateral portion of the OFC (n = 83 sites; Fig. S4). Taken together, vmPFC BGA appears tuned to the value of pleasant items, suggesting that unpleasant valuation relies on distinct mechanisms, as examined next in the aIns.

**Figure 3.**
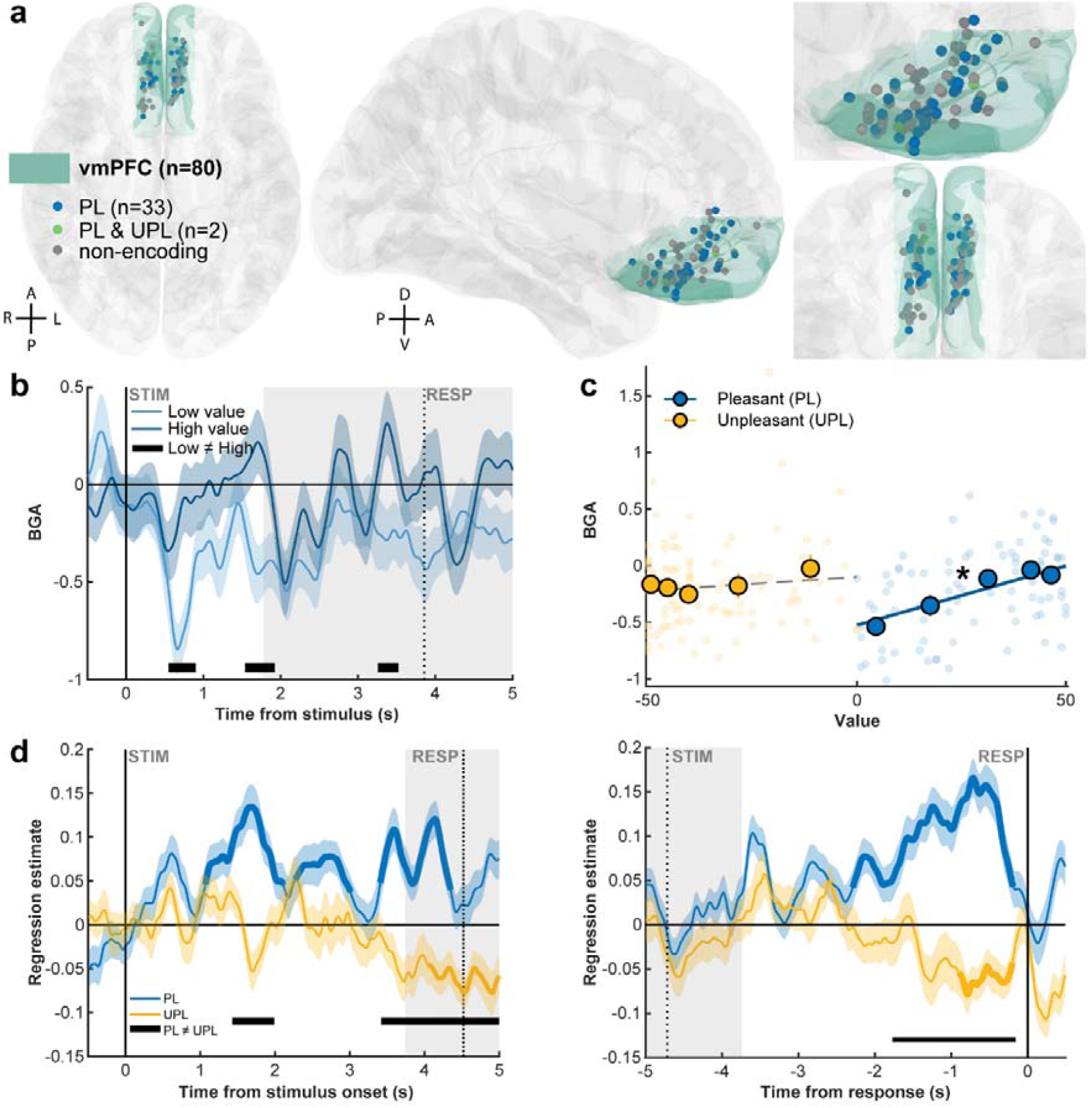
vmPFC broadband gamma activity correlates positively with the value of pleasant items. (a) Anatomical location of all 80 vmPFC recording sites from 13 patients projected onto the MNI template brain. Each site is shown as a dot and color-coded by its value-encoding profile (PL: significant correlation between BGA and pleasant value; PL & UPL: site correlated with both pleasant and unpleasant value). Anatomical orientations are indicated: anterior (A), posterior (P), dorsal (D), ventral (V), left (L), right (R). PL, pleasant; UPL, unpleasant. (b–c) Value representations from a representative vmPFC site. (b) BGA time course for low-vs. high-value trials (median split of value ratings for pleasant items). Shaded areas indicate SEM; significance is shown for illustrative purposes at uncorrected p < 0.05 (two-sided, two-sample t-tests). (c) Scatter plot of trial-averaged BGA between stimulus onset and response, plotted against item value separately for pleasant and unpleasant trials. The star indicates a significant correlation for pleasant items only (p < 0.05). (d) Group-level results across all sites. Time-resolved regression estimates of the relationship between trial-wise BGA and item value, averaged across sites and shown separately for pleasant and unpleasant items. Data are locked either to stimulus onset or to response onset. Bold lines indicate significant clusters (p_corr_ < 0.05; two-sided one-sample t-tests performed against chance for pleasant or unpleasant trials separately). Shaded areas denote ± SEM; horizontal bars mark significant clusters between pleasant and unpleasant conditions (p_corr_ < 0.05).

Consistent with this idea, during the same early window after stimulus onset, aIns BGA was significantly negatively correlated with unpleasant value (0.8–1.5 s, sum(t_166_) = -535.1, p_corr_ < α_c_; Fig. 4d; n = 167 sites, 17 subjects), whereas correlations with pleasant value were positive but not significant. The difference between pleasant and unpleasant correlations was itself significant (two-sided paired t-tests, cluster-corrected across time, 1.5–2.3 s, sum(t_166_) = 306.6, p_corr_ < α_c_; Fig. 4d). A representative aIns site encoding unpleasant value signal is illustrated in Fig. 4b–c. BGA within the first two seconds after stimulus onset was higher for ratings of the most unpleasant items compared to those rated as less unpleasant (Fig. 4b; p < 0.05). Averaging BGA from stimulus onset to response, this site showed a significant negative correlation with unpleasant, but not pleasant, value (Fig. 3c, p < 0.05). At the individual site level, value coding in the aIns was more heterogeneous than in the vmPFC, with comparable numbers of sites correlating with pleasant and unpleasant item values, and correlation signs varying more widely (see Fig. 4a). Notably, 16 of 167 aIns sites showed a significant negative correlation with unpleasant value. Together, these results indicate that the aIns rapidly and preferentially encodes unpleasant value after stimulus onset across all sites (Fig. 4d), albeit with greater site-level variability. Hence, unlike the vmPFC, aIns sites showed a more balanced distribution of correlations across pleasant and unpleasant items.

**Figure 4.**
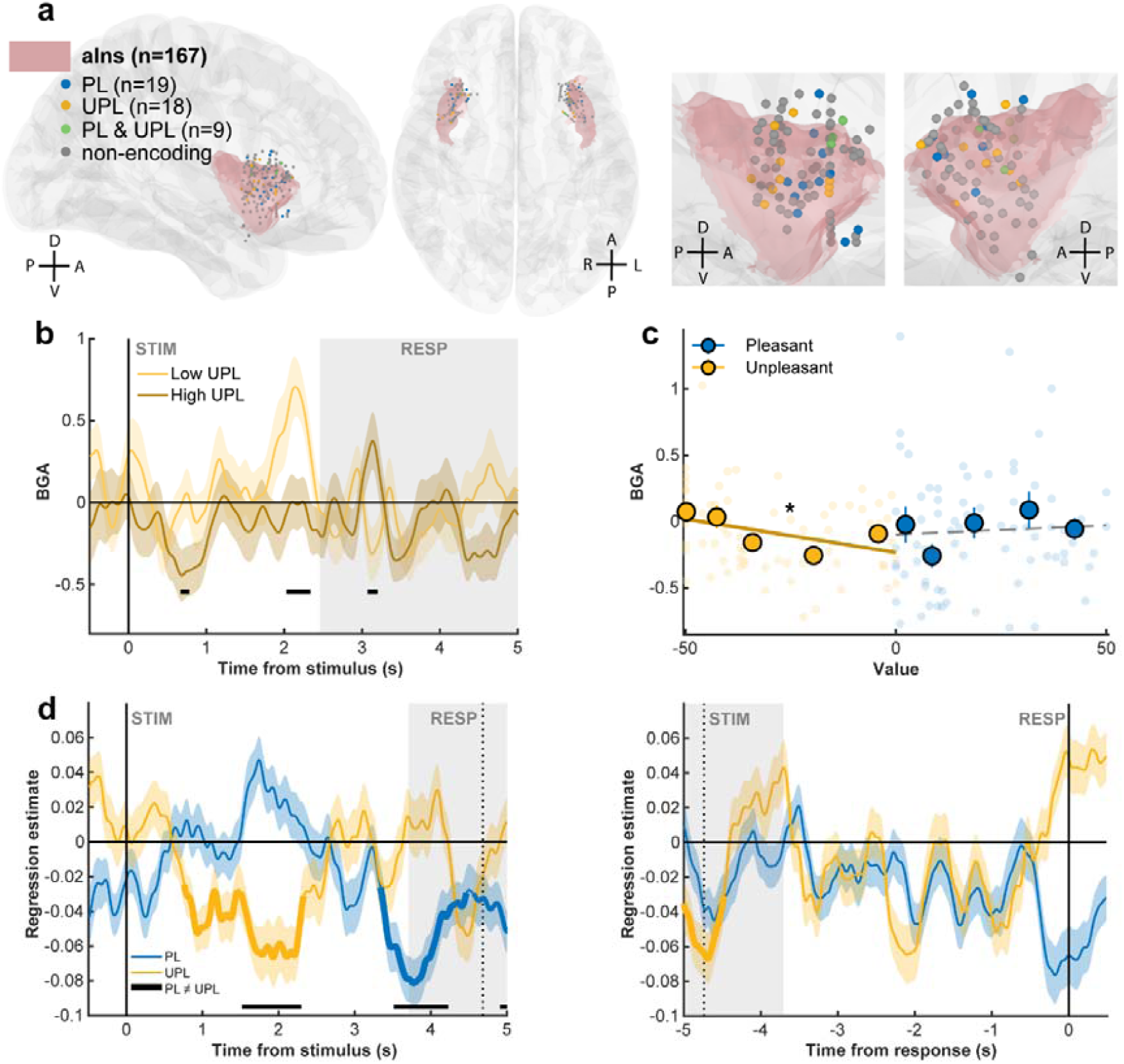
aIns broadband gamma activity correlates negatively with the value of unpleasant items. (a) Anatomical location of all 167 aIns recording sites projected onto the MNI template brain. Each site is shown as a dot and color-coded by its value-encoding profile. Anatomical orientations are indicated: anterior (A), posterior (P), dorsal (D), ventral (V), left (L), right (R). PL, pleasant; UPL, unpleasant. (b–c) Value representations from a representative aIns site. (b) BGA time course for low-vs. high-value trials (median split). Shaded areas indicate SEM; horizontal black bars mark significance between high- and low-value trials (p < 0.05, two-sided, two-sample t-tests). (c) Scatter plot of trial-averaged BGA between stimulus onset and response, plotted against item value separately for pleasant and unpleasant trials. The star indicates a significant correlation for unpleasant items only (p < 0.05). (d) Group-level results across all aIns sites. Time-resolved regression estimates of the relationship between trial-wise BGA and item value, averaged across sites and shown separately for pleasant and unpleasant items. Data are locked either to stimulus onset or to response onset. Bold lines indicate significant clusters (p_corr_ < 0.05; two-sided one sample t-tests performed against chance for pleasant or unpleasant trials separately). Shaded areas denote ± SEM; horizontal black bars mark significant clusters between pleasant and unpleasant conditions (p_corr_ < 0.05).

To further test whether opponent value-related signals in the vmPFC and aIns reflect distinct components of value, we fitted single regression models including both regions, estimated separately for pleasant and unpleasant items. To this end, we constructed all possible pairs of simultaneously recorded vmPFC and aIns sites (n = 436 pairs from 10 participants) during the rating experiment. For each pair we regressed trial-wise value against the BGA from both sites entered as separate predictors of value. Significance was assessed by testing regression estimates across pairs within each ROI at each time point (two-sided t-tests). This analysis revealed that, for pleasant items, vmPFC activity remained positively correlated with value (Fig. S5a; 1.03–5.11 s, sum(t_436_) = 3627, p_corr_ < α_c_). Similarly, for unpleasant items, aIns activity remained negatively correlated with value (Fig. S5b; peak cluster: 0.56–2.30 s sum(t_436_) = –1447, p_corr_ < α_c_). Together, these results demonstrate that the vmPFC and aIns contribute non-redundant information to value coding, beyond their known anti-correlation at rest (Kucyi et al. 2020).

We next examined whether value signals extended beyond BGA by testing correlations in other frequency ranges. Specifically, we asked whether power in distinct frequency bands correlated with the value of pleasant items in the vmPFC or with the value of unpleasant items in the aIns, focusing on sites showing the corresponding positive or negative correlations. To this end, we performed a time–frequency decomposition of sEEG signals time-locked to stimulus onset and regressed power at each time–frequency point against item value (Fig. 5). This analysis confirmed robust value-related activity in the BGA following stimulus onset in both the vmPFC (pleasant items; Fig. 5a) and aIns (unpleasant items; Fig. 5c). In the vmPFC, power across all five frequency bands significantly correlated with the value of pleasant items, showing positive correlations above 12 Hz and negative correlations below (Fig. 5b). In contrast, the aIns displayed the opposite pattern, although only low-gamma power and BGA were significant predictors of unpleasant item value (Fig. 5d). To directly compare the relative contribution of each frequency band to value coding, we developed general linear models that included power from different bands as separate regressors, testing vmPFC activity against the value of pleasant items and aIns activity against the value of unpleasant items. The key question was whether lower-frequency activity provided information beyond that carried by BGA. We therefore compared a baseline model including only BGA to all possible models combining BGA with one or more lower-frequency bands. Bayesian model selection (see Methods) identified the BGA-only model as the best account of value signals in both regions (vmPFC, pleasant value: Ef = 0.973, Xp = 1; aIns, unpleasant value: Ef = 0.945, Xp = 1). Thus, power in other frequency bands conveyed no additional information beyond BGA.

We next assessed whether and how neural activity in the vmPFC and aIns contributed to binary decisions, by examining BGA in four blocks of binary choice trials (focusing on choices between pleasant items for the vmPFC and unpleasant items for the aIns). The choice experiment included manipulations of the domain of hypothetical situations (i.e. participants had to choose between either two pleasant or two unpleasant items) and choice frame ("choose best" vs. "choose worst"). To test whether choice framing modulated value coding in the vmPFC, we examined BGA correlations with chosen and unchosen item values when participants were instructed to pick either the most or least pleasant option. In the “choose best” frame, vmPFC BGA correlated positively with the value of the chosen item after stimulus onset (Fig. 6a; pc < 0.05) and before choice onset (Fig. 6b; p < 0.05); no significant correlation was found with unchosen value. A direct comparison confirmed that BGA tracked the chosen over the unchosen value (chosen > unchosen; pc < 0.05). In the “choose worst” frame, vmPFC BGA correlated with the value of the chosen item before stimulus onset (Fig. 6c; pc < 0.05), but shifted to track the unchosen item before the response (Fig. 6d; pc < 0.05); no correlation was found for the chosen value. Thus, vmPFC activity correlated with the item that had the highest value, even when participants were instructed to select the lowest-value option. This pattern indicates that the vmPFC encodes the value of the most pleasant item available before selection, rather than the goal-congruent value (chosen value) predicted by frame-dependent accounts of value encoding in the vmPFC (Frömer et al. 2019; Castegnetti et al. 2021).

**Figure 5.**
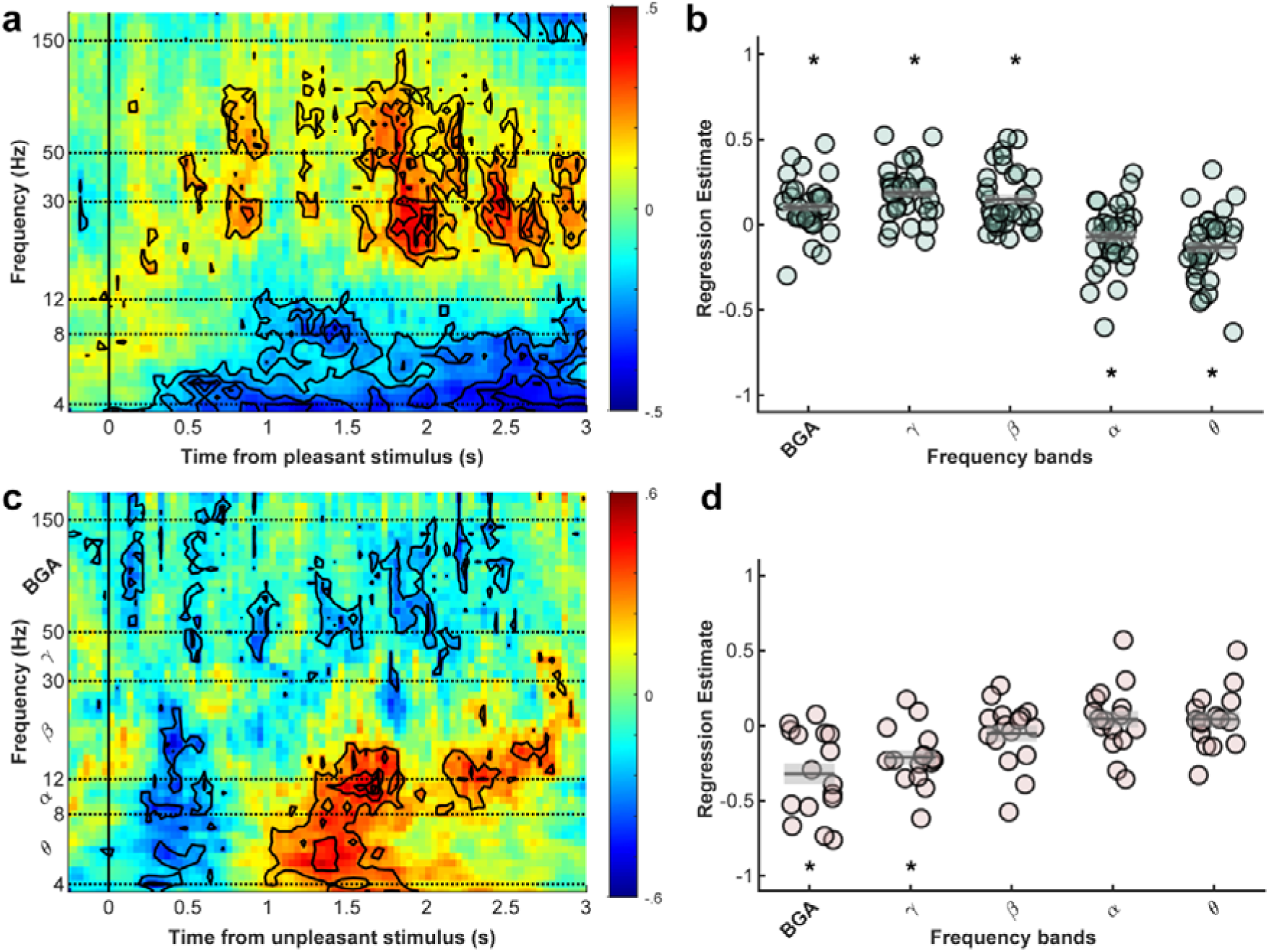
Time–frequency analysis of value-related activity across frequency bands. (a, c) Time–frequency maps of stimulus-locked regression estimates, showing correlations between power and item value, averaged across sites selectively encoding value: vmPFC sites positively correlated with pleasant value (34 sites from 10 participants) and aIns sites negatively correlated with unpleasant value (16 sites from 5 participants). Warmer colors indicate stronger positive correlations with item value. Dashed lines mark the boundaries of frequency bands that are investigated in b-d. (b, d) Regression coefficients averaged across the stimulus–response interval for each band (θ: 4–8 Hz; α: 8–12 Hz; β: 12–30 Hz; low γ: 30–50 Hz; BGA: 50–150 Hz). In the vmPFC (b), higher frequencies correlated positively and lower frequencies negatively with pleasant item value. In the aIns (d), low-γ and BGA power correlated negatively with unpleasant item value. Asterisks indicate frequency bands with regression estimates significantly different from zero (two-sided t-test, p < 0.05).

**Figure 6.**
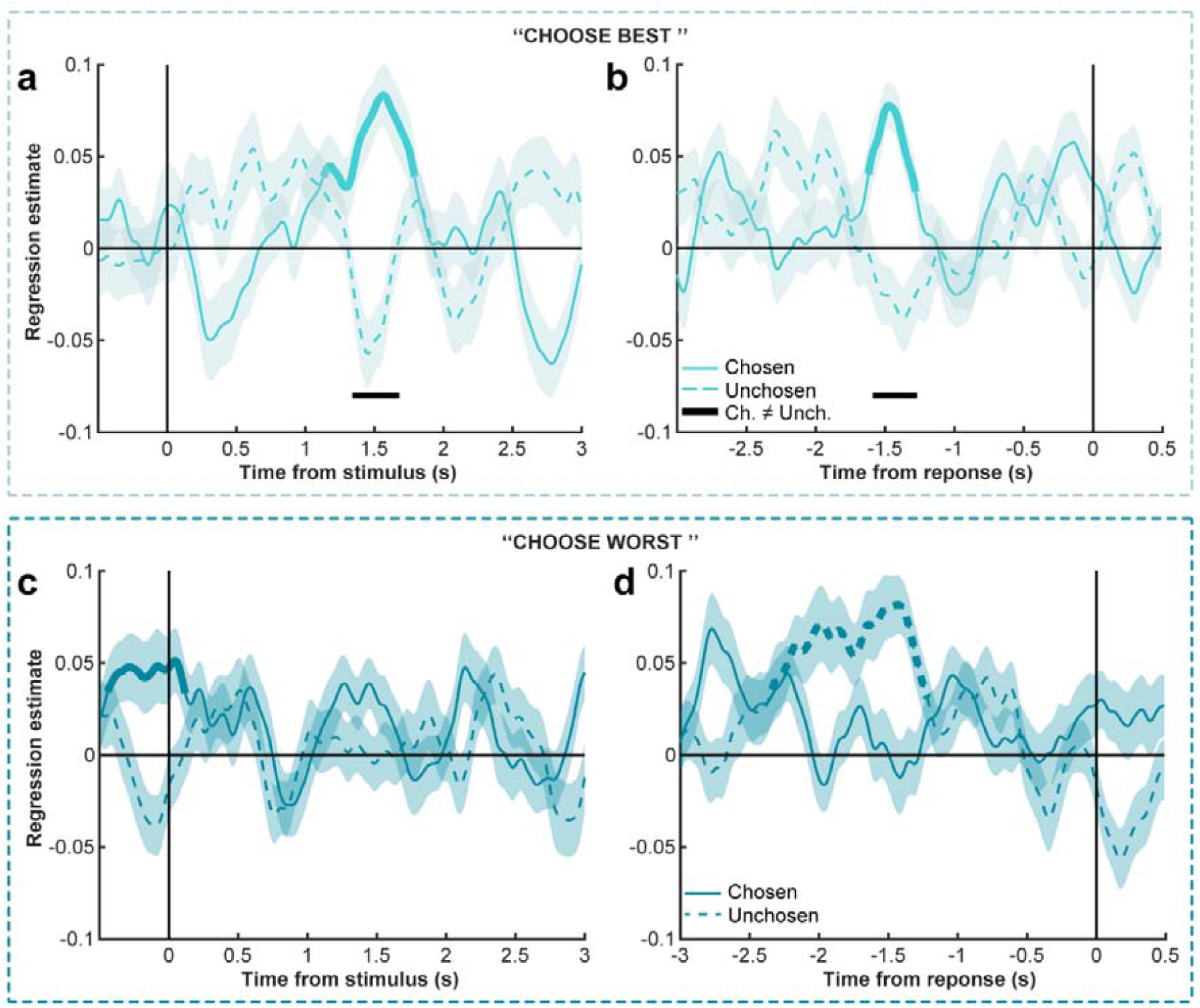
vmPFC broadband gamma activity correlates with the value of the most pleasant item. Time-resolved regression estimates of the relationship between trial-wise BGA and item value, averaged across vmPFC sites and shown separately for chosen and unchosen items in the “choose best” (a-b) and “choose worst” (c-d) conditions. Data are locked to stimulus onset (left) or response onset (right). Bold lines indicate significant clusters (p_corr_ < 0.05; two-sided one sample t-tests performed against chance for chosen or unchosen values separately). Shaded areas denote ± SEM; horizontal bars mark significant clusters between regression estimates (p_corr_ < 0.05).

We subsequently explored how choice framing influences aIns value signals. We thus computed correlations between BGA and the values of chosen and unchosen items when participants were instructed to select either the most or the least unpleasant option. When participants chose the worst option, aIns BGA correlated negatively with the chosen value after stimulus onset and before choice onset (Fig. 7c-d; pc < 0.05); no significant correlation was found with the unchosen value before the choice (although a transient positive correlation was observed at stimulus onset). Thus, when the domain (unpleasant) and choice frame (worst) were aligned, which was reflected behaviorally by faster and more consistent choices (Fig. 1f-g), aIns activity tracked the value of the most unpleasant option with a negative sign. Conversely, when participants were asked to choose the best of two unpleasant items (i.e., the more tolerable option), aIns BGA negatively tracked the value of the unchosen item from stimulus onset until just before the response, while positively encoding the chosen value prior to the response (Fig. 7a–b; pc<0.05). These results indicate that the aIns consistently signals the most aversive available option negatively before selection, even when the goal is to pick the option that is less aversive. In contrast, the chosen value reverses its sign depending on context, ruling out a mechanism purely determined by choice frame, which would produce a consistently negative association in aIns.

**Figure 7.**
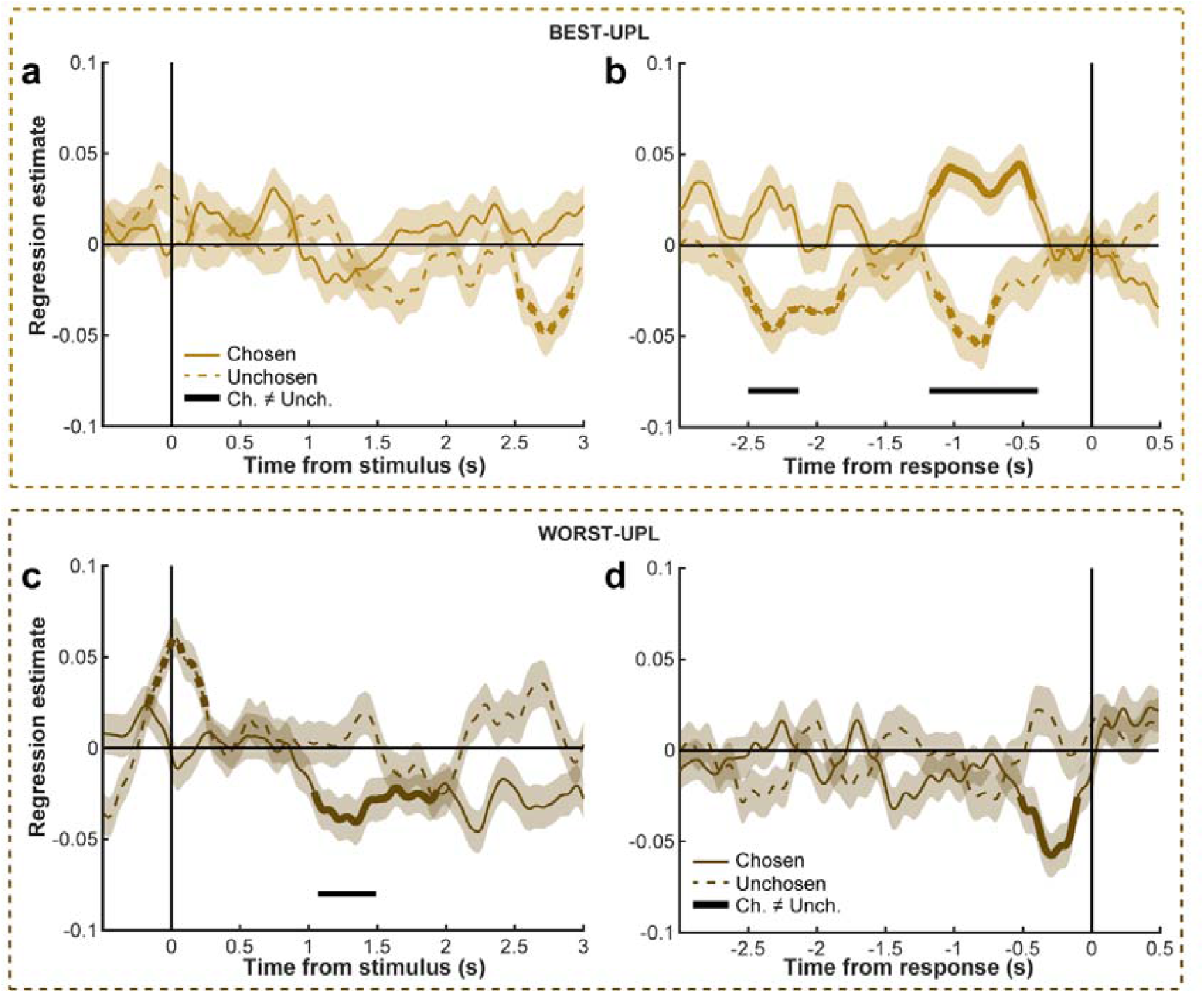
Anterior insula broadband gamma activity correlates with the value of the most unpleasant item. Time-resolved regression estimates of the relationship between trial-wise BGA and item value, averaged across aIns sites and shown separately for chosen and unchosen items in the “choose best” (a-b) and “choose worst” (c-d) conditions. Data are locked to stimulus onset (left) or response onset (right). Bold lines indicate significant clusters (p_corr_ < 0.05; two-sided one-sample t-tests performed against chance for chosen or unchosen values separately). Shaded areas denote ± SEM, horizontal bars mark significant clusters between regression estimates (p_corr_ < 0.05).

Finally, to compare the respective contributions of the vmPFC and aIns during choices, we fitted general linear models including both regions, estimated separately for each of the four choice blocks. We focused on pre-response BGA from both regions to test how both regions explained relative value (i.e., the signed difference between chosen and unchosen values).

Participants with simultaneous recordings in the vmPFC and aIns were identified (n = 10), and all possible pairs of sites were constructed (n = 389 pairs). For each pair, trial-wise relative value was regressed against BGA from both sites entered as separate predictors of relative value. For each vmPFC–aIns pair, trial-wise relative value was regressed against BGA from both sites entered as separate predictors. As shown in Fig. 8a–d, vmPFC activity correlated positively with relative value in the PL-best context and negatively in the UPL-worst context, while the opposite pattern was observed in the aIns. This sign reversal mirrors the region-specific encoding of chosen and unchosen values reported in Figs. 6–7. Visual inspection further suggested that the dynamics of relative value signals in the vmPFC and aIns were more distinguishable when item domain and choice frame were in conflict (PL-worst, UPL–best) than in more congruent conditions (PL-best or UPL-worst). To formally evaluate this observation, we assessed whether vmPFC and aIns BGA correlations with relative value diverged more during incongruent than congruent conditions using a disagreement score, defined as the difference between their time-averaged pre-response regression coefficients (in the -2.5 to 0 s time window before the response). The analysis of variance revealed a significant interaction between stimulus domain and choice frame on disagreement scores (Fig. 8e; F_(1,388)_ = 88.81, p = 4.10−19). In the pleasant domain, disagreement scores were lower in the “choose worst” condition compared with the “choose best” condition (p = 3.10−8; Tukey post-hoc test), whereas in the unpleasant domain, scores were lower in the “choose best” condition (p= 4.10−13) compared with the “choose worst” condition. These differences strikingly parallel behavioral observations we made previously: choices were less consistent with previous value ratings and reaction times longer when choice frame conflicted with the stimulus domain (Fig. 1 f-g), suggesting that the divergence in vmPFC–aIns value coding may reflect a neural mechanism underlying Pavlovian links between stimulus domain and action tendency during decision-making. To further test this idea, we checked for the existence of a correlation between disagreement scores and participants’ behavior, but none of the tested correlations reached significance (all p > 0.05), probably due to the limited statistical power in the small sample of participants with simultaneous recordings.

**Figure 8.**
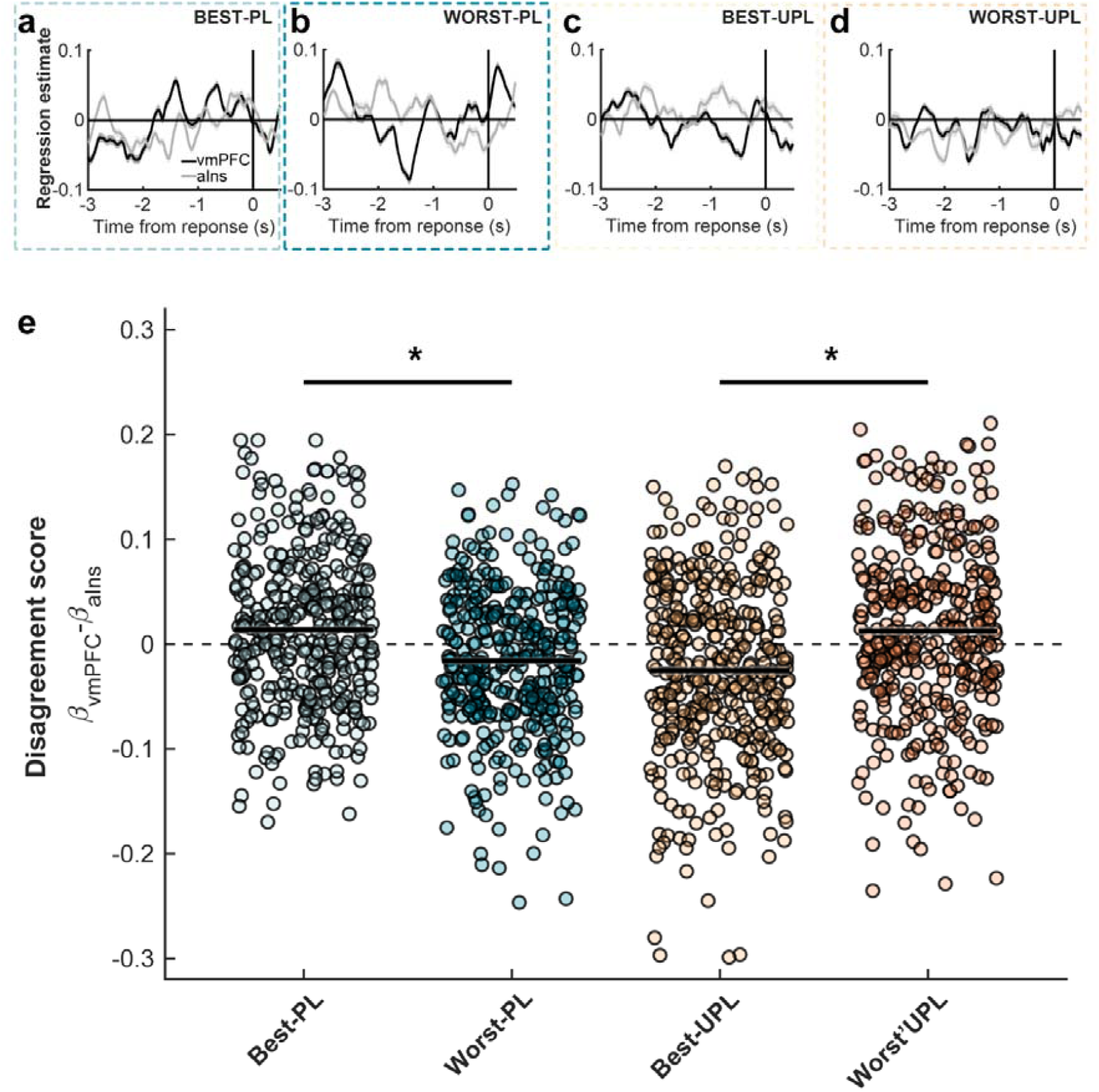
The difference between relative value signals in the vmPFC and aIns is modulated by choice context. (a-d). Time-course of regression estimates for the vmPFC and aIns BGA included together in a single GLM predicting relative value, and estimated separately for the four blocks of choices (a: Best-PL; b: Worst-PL; c: Best UPL; d: Worst UPL). Lines show the mean across all recording sites, and shaded areas indicate ± SEM across sites. (e) Disagreement scores between the vmPFC and aIns across choice contexts. Each dot represents the average difference between the vmPFC and aIns regression estimate between BGA and relative value before the response (in the -2.5 to 0 s time interval) of a given channel pair (n = 436 pairs). Asterisks indicate significance (p < 0.05). Bold horizontal lines and grey shaded areas indicate mean and SEM within each choice context. Stars indicate significance (post-hoc Tukey tests that followed the significant interaction found between stimulus domain and choice frame).

## Discussion

Intracerebral recordings acquired during a rating task (pleasant vs. unpleasant items) followed by binary decisions provided a rare opportunity to dissect the neural computations underlying value-based choice. We first characterized how the brain encodes the subjective value of hypothetical scenarios across desirable and aversive contexts. BGA correlated positively with pleasant value in the vmPFC and negatively with unpleasant value in the aIns, consistent with an anatomical dissociation between appetitive and aversive valuation circuits. During contexts in which stimulus pleasantness and choice frame were experimentally decoupled (i.e., when participants chose the least pleasant item among pleasant stimuli or the least unpleasant item among unpleasant stimuli), pre-choice BGA correlated positively with the value of the most pleasant item in the vmPFC and negatively with the value of the least unpleasant item in the anterior insula (aIns), irrespective of choice frame. The difference in correlations between BGA and relative value was larger for choices in which stimulus domain and choice frame were conflicting, consistent with slower response times and less value-driven choices in these trials. These results reveal a neural signature by which the interaction between stimulus domain and action tendencies shapes decision-making. We next discuss these findings in detail.

Using direct regression of BGA between stimulus onset and response against pleasantness ratings, we observed dissociable networks encoding value in the pleasant versus unpleasant domain. No region exhibited a domain-general value response. Instead, our findings support a functional division of labor: the vmPFC signals stimulus value in the pleasant domain (Fig. 2), while the aIns is encoding unpleasantitem value (Fig. 3). More generally, BGA increased linearly with pleasant-item value in the vmPFC, PCC and cmVC, whereas it increased as unpleasant-item value decreased in the aIns and rmVC. Crucially, the vmPFC showed robust value encoding for pleasant items, whereas neighboring OFC subregions did not. This contrasts with previous electrophysiological findings in monkeys and human intracranial EEG studies reporting similar value encoding across the vmPFC and OFC (Lopez-Persem et al. 2020; Shih et al. 2023). Exploratory inspection of OFC subregions revealed only a late negative correlation in the ventral OFC (Fig. S4). This suggests that the OFC may contribute to later, context-dependent modulation of value signals originating from the vmPFC in the context of the valuation of pleasant text items. Pleasant value encoding in the PCC was also weaker and delayed, inconsistent with its proposed role in automatic valuation (Grueschow et al. 2015). The involvement of the cmVC, encompassing parahippocampal, fusiform, and lingual gyri, is consistent with prior findings (Lopez-Persem et al. 2020), but its BGA time course was more variably related to pleasant value than in the vmPFC and PCC. Together, the latter timing and larger variability of value signals observed suggest that the pCC and cmVC contribute to value coding of pleasant items in a secondary rather than primary manner. Likewise, because the aIns was the only region showing a significant temporal cluster within the first two seconds after stimulus onset for unpleasant items, rmVC activity likely reflects later, context-dependent processing of valuation signals that originate primarily in the aIns.

### Opponent brain valuation systems for pleasant vs. unpleasant items

In the vmPFC, analyses across all recording sites and time (Fig. 2d) revealed that BGA correlated positively with the value of pleasant hypothetical situations ∼ 1 s after stimulus onset, but not with the value of unpleasant items. The correlation was consistently stronger for pleasant compared to unpleasant items. Individual site analyses further showed a highly selective response profile, with most vmPFC sites correlating only with pleasant-item value (Fig. 2a). These findings echo previous neuroimaging studies linking vmPFC activity to the valuation of neutral or pleasant stimuli (Lebreton et al. 2009, 2013; Plassmann et al. 2010; Liu et al. 2011; Bartra et al. 2013), and are consistent with additional intracerebral reports (Lopez-Persem et al. 2020; Shih et al. 2023). However, these results also challenge the notion that the vmPFC provides a domain-general value signal. Although the vmPFC can encode value across different stimulus categories (Lebreton et al. 2009; Lopez-Persem et al. 2020), here, we found much stronger signals exhibiting a consistent correlation between BGA and the value of pleasant items in the vmPFC, extending a prior observation showing that BGA in this area correlate positively and selectively with reward prediction errors but not with punishment prediction errors (Gueguen et al. 2021). Baseline vmPFC BGA also correlates positively with high, but not low, mood levels (Cecchi et al. 2021). vmPFC BGA is also reduced in response to positive self-judgments in individuals with higher depression scores (Iravani et al. 2024), indicating that automatic valuation processes in this region may be altered in the context of clinical depression. Together, these results support the hypothesis that the vmPFC is signaling the value of appetitive events.

Strikingly, the timing of this value signal remains consistent across paradigms, emerging within the .4–1.5 s range in previous visual rating and auction tasks (Lopez-Persem et al. 2020; Shih et al. 2023). In our text-based item rating paradigm, its slightly delayed and more sustained profile (1.1–2.1 s; Fig. 2d) likely reflects the additional imaginative demands of our text stimuli. In monkeys, in contrast, BGA in the OFC rapidly encodes value during economic decision tasks (∼0.2–0.4 s post-stimulus;) with shorter reaction times (< 1 s) typical of over-trained animals (Sharma et al. 2025). Yet, others have reported more sustained value-related BGA in the OFC, lasting beyond 1 s during reward expectation paradigms (Rich and Wallis 2017), suggesting that the precise neural dynamics underlying value is task-dependent. Another striking finding was that nearly all value-responsive BGA sites in the vmPFC correlated positively with the value of pleasant items (Fig. 3a), whereas only a small fraction (2 out of 35) showed negative correlations with unpleasant items. This sharply contrasts with the mixed value coding typically observed in the monkey OFC, where roughly half of the sites show either positive or negative value tuning, whether assessed through BGA or firing rates (Rich and Wallis 2017; Sharma et al. 2025). In monkeys, single neurons can also increase or decrease firing selectively for rewarding, aversive, or both types of stimuli (Morrison and Salzman 2009). In comparison, one of the few studies targeting the vmPFC in monkeys reported predominantly positive value across neurons, while OFC populations did not reliably encode value (Abitbol et al. 2015), underscoring the functional relevance of the vmPFC for valuation in monkeys. Considering the established link between BGA and local spiking activity, our results suggest that human vmPFC harbors a predominantly appetitive valuation code, driven by a larger population of neurons selectively increasing activity with the value of pleasant options.

A complementary picture emerged in the anterior insula (aIns), where BGA was negatively correlated with the value of unpleasant items. Consistent with converging evidence from fMRI (Bartra et al. 2013; Skvortsova et al. 2014; Corradi-Dell’Acqua et al. 2016; Le Bouc and Pessiglione 2022) and intracerebral recordings (Cecchi et al. 2021; Gueguen et al. 2021; Liu et al. 2021; Soyman et al. 2022), we found that aIns BGA correlated negatively with the value of unpleasant situations approximately 1 s after stimulus onset. This latency matches prior iEEG findings showing that aIns BGA and spiking activity covary with the perceived intensity of pain in a similar temporal window (Soyman et al. 2022). Unlike in the vmPFC, where value coding built up toward the response, aIns activity reflected a rapid, stimulus-locked encoding of aversive value at the population level (i.e., across all available sites), albeit with a higher variability when inspecting individual sites (Fig. 3 a-d). This pattern of results is compatible with the known contribution of this area to processing aversive and risky events. In monkeys, aIns neurons encode both reward and loss value signals (Mizuhiki et al. 2012; Yang et al. 2022), with a predominance of neurons signaling loss value—mirroring our population-level bias toward unpleasant stimuli. Human lesion and fMRI studies also highlight the aIns as a key node for risk aversion and punishment-based learning (Kuhnen and Knutson 2005; Clark et al. 2008; Palminteri et al. 2012). Together, these results position the aIns as a critical locus for encoding aversive value signals in humans, with BGA providing a temporally precise marker of this process. The timing and correlation sign of the signal suggest that the aIns contributes early stimulus-driven information about the negative (or punishing) dimension of value, complementing the vmPFC’s role in integrating appetitive value over time. Taken together, in the valuation task, we found opponent activity between the vmPFC and aIns, in line with a partially-dissociated decision-making system (Pessiglione and Delgado 2015).

During choices, our design dissociated neural signals that typically covary: those aligned with the choice frame (i.e., chosen value) and those reflecting the value of the most pleasant or most unpleasant item. The choice-frame hypothesis predicts that brain activity should scale with the difference between chosen and unchosen values, across choice frames (Hunt et al. 2012; Boorman et al. 2013; Le Bouc and Pessiglione 2022). Instead, we found that vmPFC BGA correlated positively with the most pleasant item irrespective of choice frame (Fig. 6), whereas aIns BGA showed an opposite, negative correlation with the most unpleasant item (Fig. 7). This finding contrasts with reports of vmPFC value representations being modulated by goal frame (Frömer et al. 2019; Castegnetti et al. 2021). Pre-choice activity in both the vmPFC and aIns reflected option value rather than task goal, with the vmPFC tracking the most pleasant and the aIns the most unpleasant item, irrespective of whether they had to be chosen or avoided. This suggests an intrinsic, automatic coding of value in these regions, consistent with prior fMRI (Lebreton et al. 2009) and sEEG evidence (Lopez-Persem et al. 2020) and with models framing choice as a comparison between preferred and alternative options (Lopez-Persem et al. 2016). We also observed unexpected vmPFC and aIns responses, including transient positive correlations with chosen value even before stimulus onset (Fig. 6) and positive correlations with chosen or unchosen values in the aIns (Fig. 6–7). These atypical effects may reflect a mismatch between task instructions and the brain’s habitual representation of appetitive and aversive values, leading to transient distortions in value encoding, consistent with the sign reversals previously reported for OFC activity (Lopez-Persem et al. 2020). Such deviations were most pronounced in the less natural choice contexts specifically designed to dissociate alternative forms of value coding.

At the behavioral level, reaction times were longer and choices less consistent when stimulus domain and choice frame were incongruent (Fig. 1), notably when participants had to choose the worst of pleasant scenarios or the best of unpleasant ones. This pattern aligns with the natural coupling between reward and approach, and punishment and avoidance. A similar interaction emerged at the neural level (Fig. 8): vmPFC and aIns correlations with relative value diverged more during incongruent than congruent conditions, as indexed by a disagreement score computed from their average pre-response regression coefficients for a sample of participants with simultaneous recordings. Disagreement scores were lower in the “choose worst” than “choose best” frame for pleasant items, and the reverse held for unpleasant items (Fig. 8e). These results suggest that incongruence between choice frame and stimulus domain disrupts the coherence of valuation circuits, possibly contributing to behavioral interference. However, we found no direct correlation between neural disagreement and behavioral effects across participants, likely due to limited statistical power in the subset with simultaneous recordings from both regions.

### Limitations

Although recordings were obtained from patients with drug-resistant epilepsy, multiple lines of evidence argue against contamination by pathological activity (Lopez-Persem et al. 2020; Cecchi et al. 2021; Gueguen et al. 2021). Nonlesional epileptic sites generate normal physiological responses (Liu and Parvizi 2019), and while some residual spikes or pathological high-frequency events may persist despite artifact rejection, these cannot plausibly account for the robust, value-specific BGA correlations observed across a large and heterogeneous cohort. iEEG provides unique access to the dynamics of valuation, combining millisecond precision, direct sampling of deep regions such as the vmPFC and aIns, and comparability with single-neuron recordings in monkeys. However, implantations follow clinical constraints, leading to uneven spatial coverage and limited sensitivity in unsampled or sparsely covered regions, such as the ventral striatum. Also, our task likely involved additional processes on the top of subjective valuation such as mental simulation ( induced by the use of text items), it was reassuring to observed that the vmPFC showed relatively early positive value encoding, which is more consistent with a valuation process rather than imagination, which was shown to recruit the hippocampus during a similar paradigm (Lebreton et al. 2013). Finally, while orthogonalizing choice frame and stimulus domain allowed us to disentangle several valuation processes at the neural level, such a design also introduced relatively unnatural choices in incongruent conditions.

In summary, we used human intracerebral BGA to test whether subjective value is encoded by a single uniform circuit across pleasant and unpleasant contexts. Rather than a common system, we observed dissociable circuits: vmPFC signals positively tracked pleasant items, while aIns signals negatively tracked unpleasant items. This dissociation persisted across rating and choice, including contexts where choice frame conflicted with item domain. Importantly, divergence between these circuits was most pronounced in incongruent conditions, paralleling longer reaction times and lower choice consistency observed at the behavioral level. These findings reveal complementary valuation networks supporting approach and avoidance behaviors and suggest that the human brain intrinsically encodes the most appetitive or the most aversive option value to guide our choices.

## Methods

### Patients

Thirty patients with drug-resistant epilepsy were recruited from six hospitals (Grenoble, n=14; Rennes, n=3; Toulouse, n=7; Nancy, n=1; Lyon, n=1; Prague, n=4). Participants were included if they were adults with medically refractory epilepsy and able to understand the task instructions, and were excluded if they could not complete the experimental session due to seizures, fatigue, or other clinical constraints. Three patients were excluded for failing to complete the task, leaving 27 patients (mean age 33.6 ± 1.9 years; 16 females) with usable behavioral data. One patient completed the task twice with different electrode placements, four months apart; both sessions were treated as independent. For iEEG analyses, one additional patient was excluded due to corrupted data files, resulting in 26 patients with usable intracranial recordings. All participants were native speakers or fluent in the language used at their hospital (French for the French sites, Czech for Prague) to ensure comprehension of the instructions. At the time of testing, all patients had stereotactic depth electrodes implanted for clinical purposes to localize seizure foci. Research sessions were conducted in between routine clinical procedures. Written informed consent was obtained from all participants, and the study was approved by the relevant ethics committees (IdRCB: 2017A03248-45).

### Experimental tasks

The experiment was divided into two tasks: a rating task and a binary choice task. Both were implemented on a laptop using custom-written PsychToolbox scripts running in Matlab 2019b. The tasks were performed in French or Czech dependent on patient nationality. Participants were seated comfortably in front of the screen, and responses were made using a gamepad (Logitech F310). During both tasks, each trial was preceded by a fixation cross presented on the screen for a random duration lasting between 0.5 and 1 s. No time limit was given for rating or choice trials, except during the training phases where participants were encouraged to respond faster than 7 s.

### Rating task

During the rating task, participants evaluated 240 hypothetical situations, with 120 typically pleasant and 120 typically unpleasant. To ensure a variety of options and allow comparisons between pleasant and unpleasant item sets, the trials from each category were further divided into sensory (pain/taste/smell/touch/hearing/vision – 60 items) and non-sensory situations (emotional/material/social - 60 items), with pleasant and unpleasant situations matched by sub-category (touch e.g.: petting a cat vs. having a pebble in your shoe). All items were randomized and divided into 12 blocks of 20 trials based on their main category (pleasant vs. unpleasant), and presented on the screen individually in written form. Participants rated each situation on a continuous scale from -50 to 50 using a cursor. The scale included three labels (not at all, neutral, enormously) and a block-specific instruction above it, either "I like it…" for pleasant trials or "I dislike it…" for unpleasant trials. To prevent confusion, a reminder screen preceded each block, indicating "How much would you like…?" or "How much would you dislike…?" Participants were encouraged to use the full range of the scale. The cursor was initially positioned randomly within ±25% of the midline to minimize potential motor biases. Participants completed a training session to ensure they understood the task. For analyses, ratings of unpleasant items were multiplied by -1, so that very unpleasant items corresponded to low values and very pleasant items to high values.

### Choice task

Based on the ratings obtained during the rating task, binary choices were generated and organized into four blocks of 60 trials each, following a 2 × 2 factorial design (2 stimulus domain: pleasant vs. unpleasant × 2 choice frame: choose best vs. choose worst). Each block contained 25% sensory choices primarily varying in mean overall value, 25% sensory choices varying in relative value, 25% non-sensory choices varying in mean value, and 25% non-sensory choices varying in relative value. Each item appeared twice across the task, once in a mean-value pairing and once in a relative-value pairing. Within each block, trial order was randomized, and block order was counterbalanced across participants. Before each block, participants were instructed to "choose the best" or "choose the worst" item of the two presented, drawn from either pleasant or unpleasant items according to the block type. Understanding of the block-specific instructions was verified through a short training session. During the task, pairs of items were presented in written form, one on the left and one on the right side of the screen, with a reminder of the instruction above ("Best situation" or "Worst situation"). Participants made their selections using the left and right buttons on a gamepad.

### Behavioral analyses

Prior to behavioral analyses, trials with outlier reaction times (RTs), defined as more than three standard deviations from the participant’s mean, were excluded. Group-level comparisons were performed using two-tailed paired Student’s t-tests or two-way within-subject ANOVAs. Significant ANOVA effects were followed up with pairwise comparisons using Tukey’s honestly significant difference procedure. Choice behavior was further analyzed using generalized linear mixed-effects models. Predictor variables included rating difference (the difference between left and right item ratings) and overall value (the sum of left and right ratings). Response variables were left choice (1 = left chosen, 0 = right chosen) and RT. Predictor variables were z-scored per subject and block, and RT was log-transformed to reduce skewness. All models included random intercepts and slopes per participant, with an appropriate link function (logit for binomial outcomes, identity for normal outcomes). RT models additionally included the absolute rating difference as a control variable, since choices with larger value differences are typically faster.

The fixed effects reported correspond to the following models:

Model 1: Left chosen ∼ constant + rating difference + (constant + rating difference | subject)

Model 2: RT ∼ constant + absolute rating difference + overall value + (constant + absolute rating difference + overall value | subject)

Model 3: RT ∼ constant + absolute rating difference + overall value × choice frame + (constant + absolute rating difference + overall value + choice frame | subject)

### sEEG acquisition

In all centers, the implanted sEEG electrodes were connected to a 128- or 256-channel audio-video EEG system (Micromed, Treviso, Italy) sampling at 512 or 1024 Hz. Signals were band-pass filtered online between 0.1 and 200 Hz and referenced to a common white matter electrode, except in Prague, where a Natus systems was used (Quantum NeuroWorks with a 0.01–682 Hz band-pass filter). Between 10 and 18 electrodes were implanted per patient, depending on the clinical structures of interest. Electrodes were of the Dixi type (diameter: 0.8 mm) and consisted of 8 to 18 contacts, each 2 mm in length with 1.5 mm spacing between adjacent contacts. In Toulouse, some recordings were performed using Neuralynx (ATLAS, Neuralynx, Inc.). Data were acquired using a referential montage, with the reference electrode located in the white matter. Before analysis, all signals were re-referenced to their nearest neighbor on the same electrode, yielding a bipolar montage. Intracranial EEG recordings acquired from the depth electrodes were then preprocessed using a combination of the FieldTrip toolbox (Oostenveld et al. 2011) and custom-written MATLAB scripts.

### Electrode localization and ROI definitions

Coordinates and labeling of depth electrode contacts were obtained using the semi-automatic IntrAnat Electrodes pipeline (Deman et al. 2018), implemented as a BrainVISA toolbox (Rivière et al. 2009). For each participant, a pre-implantation T1-weighted MRI scan was co-registered with either a post-implantation CT (Computed Tomography) or T1-weighted MRI scan using SPM12 (Ashburner 2009). Pre-implantation images were normalized to MNI space using SPM12 and segmented with FreeSurfer (Fischl et al. 2002). Neuroanatomical parcellations were obtained based on two atlases: the Destrieux atlas (Destrieux et al. 2010), and the MarsAtlas (Auzias et al. 2016). The Destrieux parcellation was computed with FreeSurfer. To obtain the MarsAtlas parcellation, cortical surface reconstruction was performed using the Morphologist pipeline of BrainVISA, and the final parcellation was derived using the HIP-HOP method (Auzias et al. 2016). Depth electrodes were positioned manually based on post-implantation image artifacts. Their coordinates were stored in the pre-implantation T1 “scanner-based” referential and subsequently transformed to Montreal Neurological Institute (MNI) space using the stored normalization parameters. For each contact, MarsAtlas and Destrieux anatomical labels were retrieved from the scanner-based and MNI coordinates, respectively. MarsAtlas, a cortical parcellation model specifically designed for functional data analysis, was used as the reference for labeling all cortical regions—excluding the insular cortex. In this way, 40 potential cortical ROIs were obtained per hemisphere. Given our specific interest in insular subregions, we used the Destrieux atlas instead to obtain a finer-grained parcellation of insular gyri and sulci. The obtained labels, combined with MNI coordinates, were used to segment the insula into two subregions of interest: the anterior insula, comprising the anterior (y > 5 in MNI space) portions of parcels 18 (short insular gyri), 47 (anterior circular sulcus), 48 (inferior circular sulcus), and 49 (superior circular sulcus); and the posterior insula, comprising the posterior (y < 5 in MNI space) portions of parcels 17 (central sulcus and long insular gyri), 48, and 49. Additional anatomical constraints were applied to the vmPFC to eliminate potential outlier contacts; specifically, dorsal (z = 10 in MNI space) and lateral (x = ±12 in MNI space) limits were imposed. To optimize statistical power, no distinction was made between left and right hemispheres.

### sEEG analyses

#### Analyses on broadband gamma activity

BGA extraction followed previously established procedures (Lachaux et al. 2012; Gueguen et al. 2021; Cecchi et al. 2022). Signals from each channel were re-referenced to their nearest neighbor, yielding a bipolar montage, and recordings sampled at 1024 Hz were downsampled to 512 Hz. Each bipolar signal was then band-pass filtered in successive 10-Hz frequency bands from 50 to 150 Hz using a non-causal zero-phase finite impulse response filter (roll-off: 0.5 Hz). For each filtered signal, the envelope was computed via the Hilbert transform and normalized across the full recording session by dividing by its mean and multiplying by 100, yielding a time series expressed as a percentage of the mean. Envelopes from the 10 sub-bands were averaged to obtain a single BGA time series per bipole. These time series were then smoothed with a 250-ms sliding window, downsampled to 100 Hz (one sample every 10 ms), and epoched relative to stimulus onset (-1 to 6 s) and response onset (-6 to 1 s).

Artifacts were identified and rejected on a trial-by-trial basis per channel to minimize data loss. Trials with reaction times >20 s were excluded. Trials showing outlier mean or maximum BGA values (>3 SD from the channel mean) were also removed. Channels exhibiting inconsistent activity were identified based on the standard deviation of their remaining trials’ mean and maximum BGA; those exceeding one SD from the overall mean (computed across all channels) were excluded. Data were then visually inspected, and two patients were discarded due to generalized artifacts across channels and trials that could not be corrected by outlier removal. This procedure was applied separately for the rating and choice tasks. The final dataset retained 83.3% of channels and 91.5% of trials from the rating task, and 81.4% of channels and 90.54% of trials from the choice task. All remaining trials were baseline-corrected using the mean activity from -250 to -50 ms before stimulus onset, averaged across all trials.

### General linear models (GLMs)

For all analyses, regressors were z-scored to allow comparability of regression coefficients across participants and variables. When several regressors were included in a single model, multicollinearity was assessed by calculating the variance inflation factor (Belsley et al. 1980), with a maximum upper limit set to 3.

#### sEEG analyses of the rating task

In a first two GLMs were designed to identify brain regions encoding value (two separate GLMs for the value of pleasant or unpleasant items were tested), single-trial, time-averaged BGA power (*P*), computed between stimulus onset and response onset and was regressed across trials against item value (V):

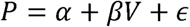

With α corresponding to the intercept, β corresponding to the regression estimate on which statistical tests are conducted and ε corresponding to the error term.

For this pseudo–whole-brain approach, the resulting p-values obtained for each t ROI were corrected for multiple comparisons across ROIs using the Bonferroni method. Within each ROI, a site was considered to significantly encode value if BGA within this time window significantly correlated with value across trials. The same two GLMs were used for the time-resolved analyses, except that BGA was regressed across trials against value at every time point, and therefore, significance was assessed using permutation tests which aimed at correcting for multiple comparison in time and ROIs dimensions (see below).

Next, to further assess whether vmPFC and aIns regions would represent separate components of value, value was regressed across trials against BGA of these two regions at every time point as follows (two separate GLMs were tested for the value of pleasant or unpleasant items):

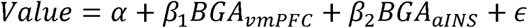

With α corresponding to the intercept, {3_1_ and {3_2_ corresponding to the regression estimates on which statistical tests are conducted and ε corresponding to the error term. For each participant with sites in both ROIs, regression was done for all possible pairs of vmPFC and aIns recording sites recorded within a given participant.

To investigate the contribution of lower frequency bands, time–frequency analyses were carried out using the FieldTrip toolbox for Matlab. Spectral powers were estimated using a ‘multitapering’ time–frequency transform (Slepian tapers, lower frequency range:4–32 Hz, 6 cycles and 3 tapers per window; higher frequency range: 32–200 Hz, fixed time windows of 187.5 ms, 4–31 tapers per window).

This approach uses a constant number of cycles across frequencies up to 32 Hz (hence a time window with a duration that decreases when frequency increases), and a fixed time window with an increasing number of tapers above 32 Hz, to obtain more precise power estimates by adaptively increasing smoothing at high frequencies. To assess the contribution of the different frequency bands to value representation, value was regressed across trials against power P (normalized envelope) of each frequency band, averaged over time between – stimulus onset to choice onset:

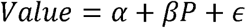

With α corresponding to the intercept and ε to the error term. The significance of the regression estimates β was assessed across recording sites using two-sided, one-sample, Student’s t-tests.

In order to determine whether other frequency bands provided additional information relative to the BGA, the following sixteen GLMs were compared:

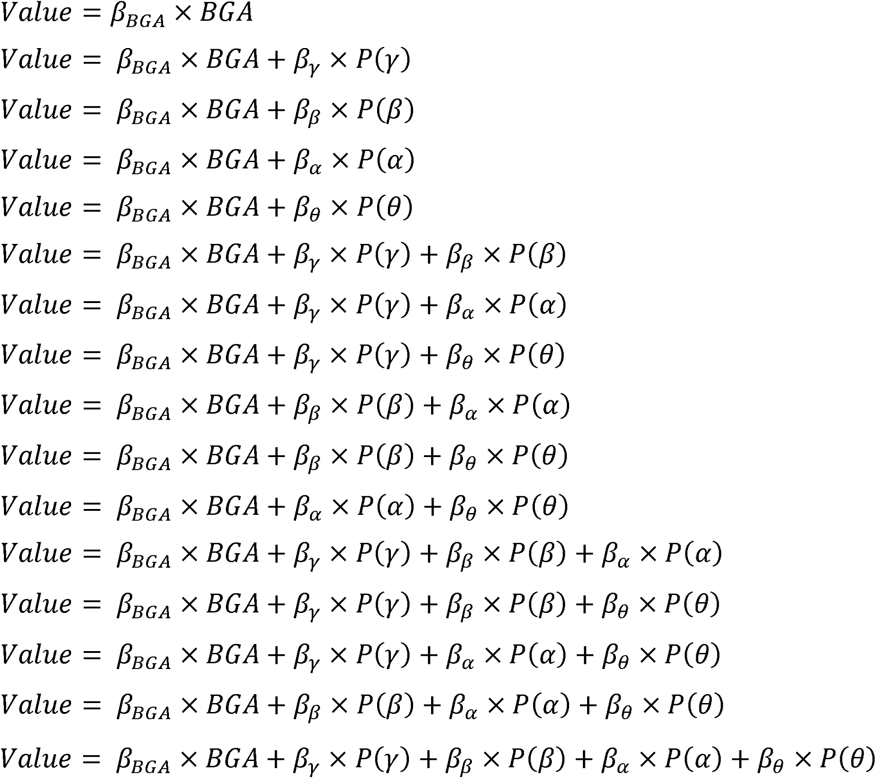

With β corresponding to the regression estimates, and P each power time series averaged between stimulus to response onset in five distinct frequency bands (θ: 4–8 Hz; α: 8–12 Hz; β: 12–30 Hz; γ: 30–50 Hz; BGA: 50–150 Hz).

The model comparison was conducted using the VBA toolbox (Daunizeau et al. 2014). Log-model evidence obtained in each recording site was taken to a group-level, random-effect, Bayesian model selection (RFX-BMS) procedure (Rigoux et al. 2014). RFX-BMS provides an exceedance probability that measures how likely it is that a given model is more frequently implemented, relative to all the others considered in the model space, in the population from which samples were drawn.

#### sEEG analyses of the choice task

To test how value signals were expressed across choice contexts, BGA was regressed across trials against chosen and unchosen value at every time point (four GLMs tested separately for the 2 domain by two choice frame choice contexts):

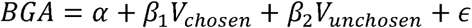

With α corresponding to the intercept, {3_1_ and {3_2_ corresponding to the regression estimates on which statistical tests are conducted and ε corresponding to the error term.

To compare the contributions of the vmPFC and aIns in explaining relative value during choices, we fitted general linear models including both regions, estimated separately for each of the four choice blocks. For each of these four GLMs, relative value was regressed across trials against BGA of these two regions at every time point as follows:

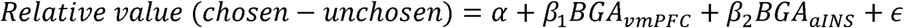

With α corresponding to the intercept, {3_1_ and {3_2_ corresponding to the regression estimates on which statistical tests are conducted and ε corresponding to the error term. For each participant with sites in both ROIs, regression was done for all possible pairs of vmPFC and aIns recording sites recorded within a given participant.

Finally, a disagreement score between vmPFC and aINS was estimated for each pair of vmPFC-aIns sites, by first averaging coefficients within the -2.5 to 0 s time window pre-response and then computing the difference between vmPFC (β_1_) and daIns (β_2_) coefficients. Disagreement scores were compared across conditions using a two-way within-participants ANOVA, followed by pairwise comparisons (using Tukey’s honestly significant difference procedure).

### Permutation tests

For all time-resolved GLMs, we used a non-parametric, cluster-based permutation procedure to correct for multiple comparisons across time. First, for each site, the correspondence between trial-wise regressors and neural activity was shuffled 300 times, generating site-specific null distributions of regression estimates time course. Second, for each ROI, one shuffled time course per site was randomly drawn from these 300 permutations, and this process was repeated 60,000 times to create ROI-level null distributions. For each of the 60 000 ROI-level permutation, the maximal cluster-level statistic (the maximal sum of t-values over contiguous time points exceeding a significance threshold of 0.05) was computed to generate a null distribution of effect sizes at the ROI level. This null distribution was then used to determine whether a given cluster in the original (non-shuffled) data was significant at a cluster-level corrected p-value (pcorr) ≤ 0.05 (as previously describe in Domenech et al., 2020; Gueguen et al., 2021; Cecchi et al., 2022). This statistical framework follows the principles of nonparametric cluster-based permutation testing (Maris & Oostenveld, 2007).

## Data availability

The data generated in this study will be will be made available by the corresponding author upon reasonable request.

## Code availability

All Matlab code necessary to reproduce our analyses will be made available by the corresponding author upon reasonable request.

## Acknowledgements

J.B. is supported by Agence National de la Recherche (EPICOG: ANR-22-CE17-0057) and Fondation pour la recherche medicale (FRM). C.B., J.B and G.B. are supported by ANR (ANR-15-IDEX-0002, IRS2019). JH was supported by ERDF-Project Brain dynamics, No. CZ.02.01.01/00/22_008/0004643. JH and PM are members of the Epilepsy Research Centre Prague - EpiReC consortium. Motol University Hospital is a full member of ERN EpiCARE.

## Author contributions Statement

J.B. and C.B. conceived the study. L.M., P.K., J.H., P.M., A.N., S.R., L.M., M.D. and E.B. participated in patient recruitment and inclusion. C.B., B.C.F., G.B. and J.B conducted data curation and analyses. C.B., A.L.P., M.P., and J.B. wrote the manuscript. All authors critically assessed and discussed the results, and revised and approved the manuscript at all stages.

## References

1. Abitbol R, Lebreton M, Hollard G, Richmond BJ, Bouret S, Pessiglione M. 2015. Neural Mechanisms Underlying Contextual Dependency of Subjective Values: Converging Evidence from Monkeys and Humans. J Neurosci. 35:2308–2320.

2. Ashburner J. 2009. Computational anatomy with the SPM software. Magn Reson Imaging. 27:1163–1174.

3. Auzias G, Coulon O, Brovelli A. 2016. *MarsAtlas*LJ: A cortical parcellation atlas for functional mapping:MarsAtlas. Hum Brain Mapp. 37:1573–1592.

4. Bartra O, McGuire JT, Kable JW. 2013. The valuation system: A coordinate-based meta-analysis of BOLD fMRI experiments examining neural correlates of subjective value. NeuroImage. 76:412–427.

5. Belsley DA, Kuh E, Welsch RE. 1980. Regression Diagnostics: Identifying Influential Data and Sources of Collinearity. 1st ed, Wiley Series in Probability and Statistics. Wiley.

6. Boorman ED, Rushworth MF, Behrens TE. 2013. Ventromedial Prefrontal and Anterior Cingulate Cortex Adopt Choice and Default Reference Frames during Sequential Multi-Alternative Choice. J Neurosci. 33:2242–2253.

7. Boureau Y-L, Dayan P. 2011. Opponency Revisited: Competition and Cooperation Between Dopamine and Serotonin. Neuropsychopharmacology. 36:74–97.

8. Castegnetti G, Zurita M, De Martino B. 2021. How usefulness shapes neural representations during goal-directed behavior. Sci Adv. 7:eabd5363.

9. Cecchi R, Vinckier F, Hammer J, Marusic P, Nica A, Rheims S, Trebuchon A, Barbeau E, Denuelle M, Maillard L, Minotti L, Kahane P, Pessiglione M, Bastin J. 2021. Intracerebral mechanisms explaining the impact of incidental feedback on mood state and risky choice (preprint). Neuroscience.

10. Cecchi R, Vinckier F, Hammer J, Marusic P, Nica A, Rheims S, Trebuchon A, Barbeau EJ, Denuelle M, Maillard L, Minotti L, Kahane P, Pessiglione M, Bastin J. 2022. Intracerebral mechanisms explaining the impact of incidental feedback on mood state and risky choice. eLife. 11:e72440.

11. Clairis N, Pessiglione M. 2022. Value, Confidence, Deliberation: A Functional Partition of the Medial Prefrontal Cortex Demonstrated across Rating and Choice Tasks. J Neurosci. 42:5580–5592.

12. Clark L, Bechara A, Damasio H, Aitken MRF, Sahakian BJ, Robbins TW. 2008. Differential effects of insular and ventromedial prefrontal cortex lesions on risky decision-making. Brain. 131:1311–1322.

13. Clithero JA, Rangel A. 2014. Informatic parcellation of the network involved in the computation of subjective value. Soc Cogn Affect Neurosci. 9:1289–1302.

14. Corradi-Dell’Acqua C, Tusche A, Vuilleumier P, Singer T. 2016. Cross-modal representations of first-hand and vicarious pain, disgust and fairness in insular and cingulate cortex. Nat Commun. 7:10904.

15. Daunizeau J, Adam V, Rigoux L. 2014. VBA: A Probabilistic Treatment of Nonlinear Models for Neurobiological and Behavioural Data. PLoS Comput Biol. 10:e1003441.

16. Deman P, Bhattacharjee M, Tadel F, Job A-S, Rivière D, Cointepas Y, Kahane P, David O. 2018. IntrAnat Electrodes: A Free Database and Visualization Software for Intracranial Electroencephalographic Data Processed for Case and Group Studies. Front Neuroinformatics. 12.

17. Destrieux C, Fischl B, Dale A, Halgren E. 2010. Automatic parcellation of human cortical gyri and sulci using standard anatomical nomenclature. NeuroImage. 53:1–15.

18. Fischl B, Salat DH, Busa E, Albert M, Dieterich M, Haselgrove C, Van Der Kouwe A, Killiany R, Kennedy D, Klaveness S, Montillo A, Makris N, Rosen B, Dale AM. 2002. Whole Brain Segmentation. Neuron. 33:341–355.

19. Frömer R, Dean Wolf CK, Shenhav A. 2019. Goal congruency dominates reward value in accounting for behavioral and neural correlates of value-based decision-making. Nat Commun. 10:4926.

20. Grueschow M, Polania R, Hare TA, Ruff CC. 2015. Automatic versus Choice-Dependent Value Representations in the Human Brain. Neuron. 85:874–885.

21. Gueguen MCM, Lopez-Persem A, Billeke P, Lachaux J-P, Rheims S, Kahane P, Minotti L, David O, Pessiglione M, Bastin J. 2021. Anatomical dissociation of intracerebral signals for reward and punishment prediction errors in humans. Nat Commun. 12:3344.

22. Hosokawa T, Kato K, Inoue M, Mikami A. 2007. Neurons in the macaque orbitofrontal cortex code relative preference of both rewarding and aversive outcomes. Neurosci Res. 57:434–445.

23. Hunt LT, Kolling N, Soltani A, Woolrich MW, Rushworth MFS, Behrens TEJ. 2012. Mechanisms underlying cortical activity during value-guided choice. Nat Neurosci. 15:470–476.

24. Iravani B, Kaboodvand N, Stieger JR, Liang EY, Lusk Z, Fransson P, Deutsch GK, Gotlib IH, Parvizi J. 2024. Intracranial Recordings of the Human Orbitofrontal Cortical Activity during Self-Referential Episodic and Valenced Self-Judgments. J Neurosci. 44:e1634232024.

25. Kable JW, Glimcher PW. 2007. The neural correlates of subjective value during intertemporal choice. Nat Neurosci. 10:1625–1633.

26. Kucyi A, Daitch A, Raccah O, Zhao B, Zhang C, Esterman M, Zeineh M, Halpern CH, Zhang K, Zhang J, Parvizi J. 2020. Electrophysiological dynamics of antagonistic brain networks reflect attentional fluctuations. Nat Commun. 11:325.

27. Kuhnen CM, Knutson B. 2005. The Neural Basis of Financial Risk Taking. Neuron. 47:763–770.

28. Lachaux J-P, Axmacher N, Mormann F, Halgren E, Crone NE. 2012. High-frequency neural activity and human cognition: Past, present and possible future of intracranial EEG research. Prog Neurobiol. 98:279–301.

29. Lachaux J-P, Fonlupt P, Kahane P, Minotti L, Hoffmann D, Bertrand O, Baciu M. 2007. Relationship between task-related gamma oscillations and BOLD signal: New insights from combined fMRI and intracranial EEG. Hum Brain Mapp. 28:1368–1375.

30. Le Bouc R, Pessiglione M. 2022. A neuro-computational account of procrastination behavior. Nat Commun. 13:5639.

31. Lebreton M, Bertoux M, Boutet C, Lehericy S, Dubois B, Fossati P, Pessiglione M. 2013. A critical role for the hippocampus in the valuation of imagined outcomes. PLoS Biol. 11:e1001684.

32. Lebreton M, Jorge S, Michel V, Thirion B, Pessiglione M. 2009. An Automatic Valuation System in the Human Brain: Evidence from Functional Neuroimaging. Neuron. 64:431–439.

33. Lindquist KA, Satpute AB, Wager TD, Weber J, Barrett LF. 2016. The Brain Basis of Positive and Negative Affect: Evidence from a Meta-Analysis of the Human Neuroimaging Literature. Cereb Cortex. 26:1910–1922.

34. Liu C-C, Moosa S, Quigg M, Elias WJ. 2021. Anterior insula stimulation increases pain threshold in humans: a pilot study. J Neurosurg. 135:1487–1492.

35. Liu S, Parvizi J. 2019. Cognitive refractory state caused by spontaneous epileptic high-frequency oscillations in the human brain. Sci Transl Med. 11:eaax7830.

36. Liu X, Hairston J, Schrier M, Fan J. 2011. Common and distinct networks underlying reward valence and processing stages: A meta-analysis of functional neuroimaging studies. Neurosci Biobehav Rev. 35:1219–1236.

37. Lopez-Persem A, Bastin J, Petton M, Abitbol R, Lehongre K, Adam C, Navarro V, Rheims S, Kahane P, Domenech P, Pessiglione M. 2020. Four core properties of the human brain valuation system demonstrated in intracranial signals. Nat Neurosci.

38. Lopez-Persem A, Domenech P, Pessiglione M. 2016. How prior preferences determine decision-making frames and biases in the human brain. eLife. 5.

39. Manning JR, Jacobs J, Fried I, Kahana MJ. 2009. Broadband Shifts in Local Field Potential Power Spectra Are Correlated with Single-Neuron Spiking in Humans. J Neurosci. 29:13613–13620.

40. Mizuhiki T, Richmond BJ, Shidara M. 2012. Encoding of reward expectation by monkey anterior insular neurons. J Neurophysiol. 107:2996–3007.

41. Morrison SE, Salzman CD. 2009. The Convergence of Information about Rewarding and Aversive Stimuli in Single Neurons. J Neurosci. 29:11471–11483.

42. Mukamel R, Gelbard H, Arieli A, Hasson U, Fried I, Malach R. 2005. Coupling between neuronal firing, field potentials, and FMRI in human auditory cortex. Science. 309:951–954.

43. Nicolle A, Klein-Flügge MC, Hunt LT, Vlaev I, Dolan RJ, Behrens TEJ. 2012. An Agent Independent Axis for Executed and Modeled Choice in Medial Prefrontal Cortex. Neuron. 75:1114–1121.

44. Niessing J. 2005. Hemodynamic Signals Correlate Tightly with Synchronized Gamma Oscillations. Science. 309:948–951.

45. Nir Y, Fisch L, Mukamel R, Gelbard-Sagiv H, Arieli A, Fried I, Malach R. 2007. Coupling between Neuronal Firing Rate, Gamma LFP, and BOLD fMRI Is Related to Interneuronal Correlations. Curr Biol. 17:1275–1285.

46. Oostenveld R, Fries P, Maris E, Schoffelen J-M. 2011. FieldTrip: Open source software for advanced analysis of MEG, EEG, and invasive electrophysiological data. Comput Intell Neurosci. 2011:156869.

47. Palminteri S, Justo D, Jauffret C, Pavlicek B, Dauta A, Delmaire C, Czernecki V, Karachi C, Capelle L, Durr A, Pessiglione M. 2012. Critical Roles for Anterior Insula and Dorsal Striatum in Punishment-Based Avoidance Learning. Neuron. 76:998–1009.

48. Pessiglione M, Delgado MR. 2015. The good, the bad and the brain: neural correlates of appetitive and aversive values underlying decision making. Curr Opin Behav Sci. 5:78–84.

49. Plassmann H, O’Doherty JP, Rangel A. 2010. Appetitive and Aversive Goal Values Are Encoded in the Medial Orbitofrontal Cortex at the Time of Decision Making. J Neurosci. 30:10799–10808.

50. Rich EL, Wallis JD. 2017. Spatiotemporal dynamics of information encoding revealed in orbitofrontal high-gamma. Nat Commun. 8:1139.

51. Rigoux L, Stephan KE, Friston KJ, Daunizeau J. 2014. Bayesian model selection for group studies — Revisited. NeuroImage. 84:971–985.

52. Rivière D, Geffroy D, Denghien I, Souedet N, Cointepas Y. 2009. BrainVISA: an extensible software environment for sharing multimodal neuroimaging data and processing tools. NeuroImage. 47:S163.

53. Sepulveda P, Usher M, Davies N, Benson AA, Ortoleva P, De Martino B. 2020. Visual attention modulates the integration of goal-relevant evidence and not value. eLife. 9:e60705.

54. Sharma D, Lupkin SM, McGinty VB. 2025. Orbitofrontal High-Gamma Reflects Spike-Dissociable Value and Decision Mechanisms. J Neurosci. 45:e0789242025.

55. Shih W-Y, Yu H-Y, Lee C-C, Chou C-C, Chen C, Glimcher PW, Wu S-W. 2023. Electrophysiological population dynamics reveal context dependencies during decision making in human frontal cortex. Nat Commun. 14:7821.

56. Skvortsova V, Palminteri S, Pessiglione M. 2014. Learning To Minimize Efforts versus Maximizing Rewards: Computational Principles and Neural Correlates. J Neurosci. 34:15621–15630.

57. Soyman E, Bruls R, Ioumpa K, Müller-Pinzler L, Gallo S, Qin C, Van Straaten EC, Self MW, Peters JC, Possel JK, Onuki Y, Baayen JC, Idema S, Keysers C, Gazzola V. 2022. Intracranial human recordings reveal association between neural activity and perceived intensity for the pain of others in the insula. eLife. 11:e75197.

58. Tang H, Bartolo R, Averbeck BB. 2024. Ventral frontostriatal circuitry mediates the computation of reinforcement from symbolic gains and losses. Neuron. 112:3782–3795.e5.

59. Tom SM, Fox CR, Trepel C, Poldrack RA. 2007. The neural basis of loss aversion in decision-making under risk. Science. 315:515–518.

60. Yang Y-P, Li X, Stuphorn V. 2022. Primate anterior insular cortex represents economic decision variables proposed by prospect theory. Nat Commun. 13:717.

