## Supplemental figures and tables for "Choosing the best or avoiding the worst: complementary and opponent value signals in the human brain"

### Supplementary materials

#### Supplementary figures

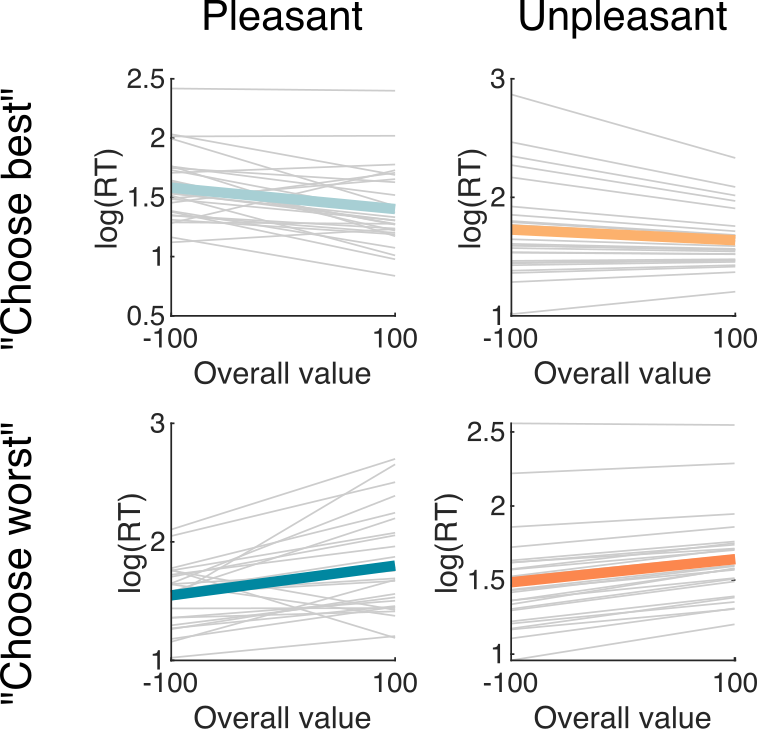

Figure S1. Overall value effects on reaction time depend on task goal. Generalized mixed-effects models were applied to predict RT from the summed value of presented options. Colored lines represent fixed effects while grey lines represent random effects (one line per subject). Overall value was only significantly associated with RT when participants were asked to choose the worst from two pleasant options (β=0.03±0.01, t_(1573)_=2.02, p=0.044) or from two unpleasant options (β=0.02±0.01, t_(1606)_=2.17, p=0.03). However, aggregating "choose best" and "choose worst" data revealed a significant interaction between overall value and task goal for both pleasant (β=-0.05±0.01, t_(3176)_=-3.38, p=7.10^-04^) and unpleasant (β=-0.03±0.01, t_(3187)_=-2.03, p=0.042) conditions.

.

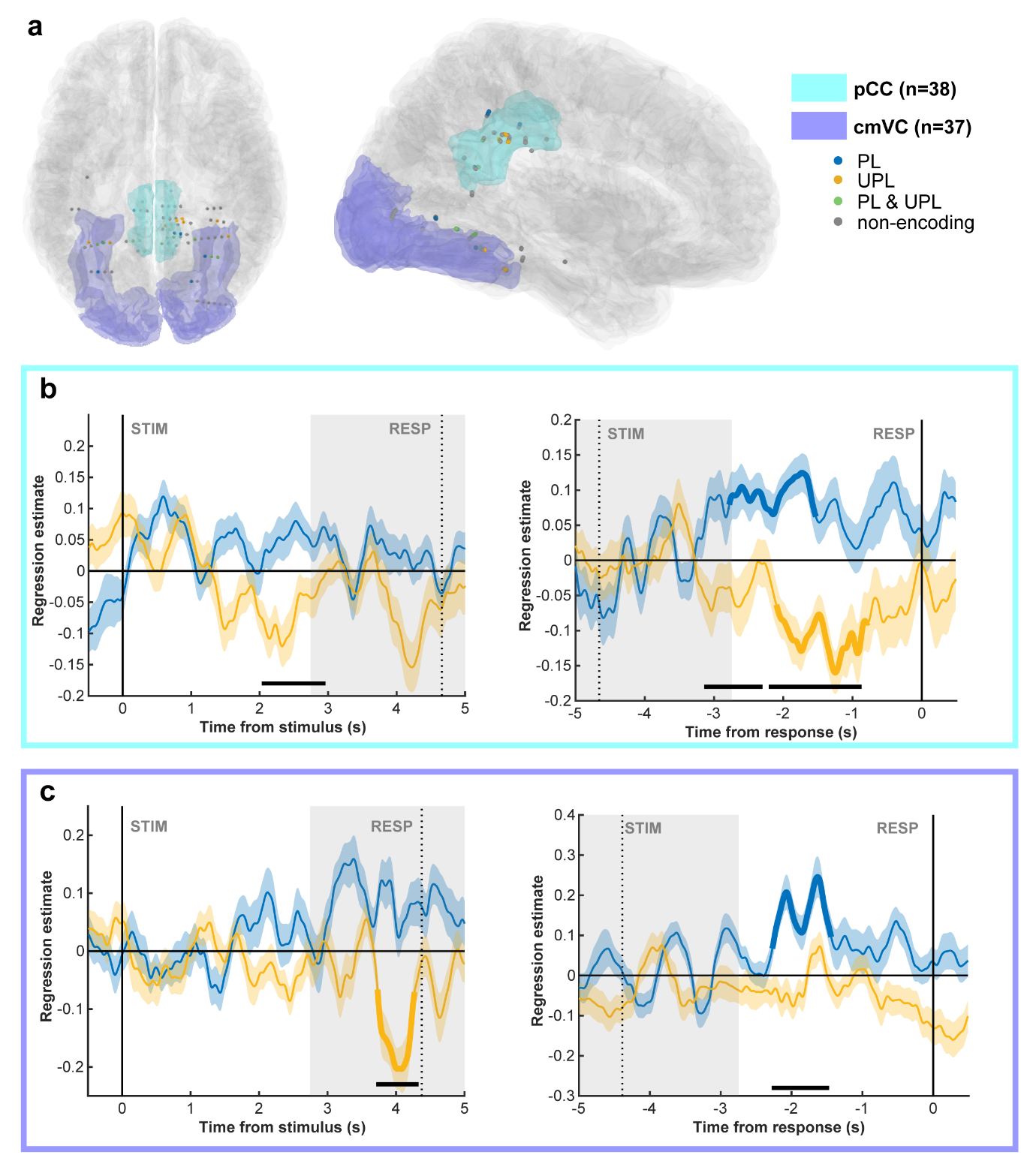

**Figure S2. Delayed positive relationship between BGA and pleasant value in the posterior cingulate and caudo-medial visual cortex.** (a) Anatomical locations of all pCC and cmVC recording sites projected onto the MNI template brain. Each site is shown as a dot, color-coded by its value-encoding profile (pCC: 38 sites; 4 PL, 4 UPL, 4 PL&UPL; cmVC: 37 sites; 4 PL, 4 UPL, 4 PL&UPL). (b–c) Group-level, time-resolved regression results across all sites for the pCC (b) and cmVC (c). Lines show regression estimates relating trial-wise BGA to item value, averaged across sites and plotted separately for pleasant and unpleasant items. Data are time-locked to stimulus onset (left panels) or response onset (right panels). Bold lines indicate significant clusters (p_corr_ < 0.05), shaded areas represent ± SEM across sites, and horizontal bars mark significant clusters ( p_corr_ < 0.05). Compared with the vmPFC (main Fig. 3), these regions showed later-onset value signals in the stimulus-locked analysis (> 5 s; panels b–c), greater variability across sites, and fewer significant PL sites (panel a; n = 4 PL per region). Additionally, the cmVC exhibited a transient negative correlation with unpleasant value in the response-locked analysis (right panel of c).

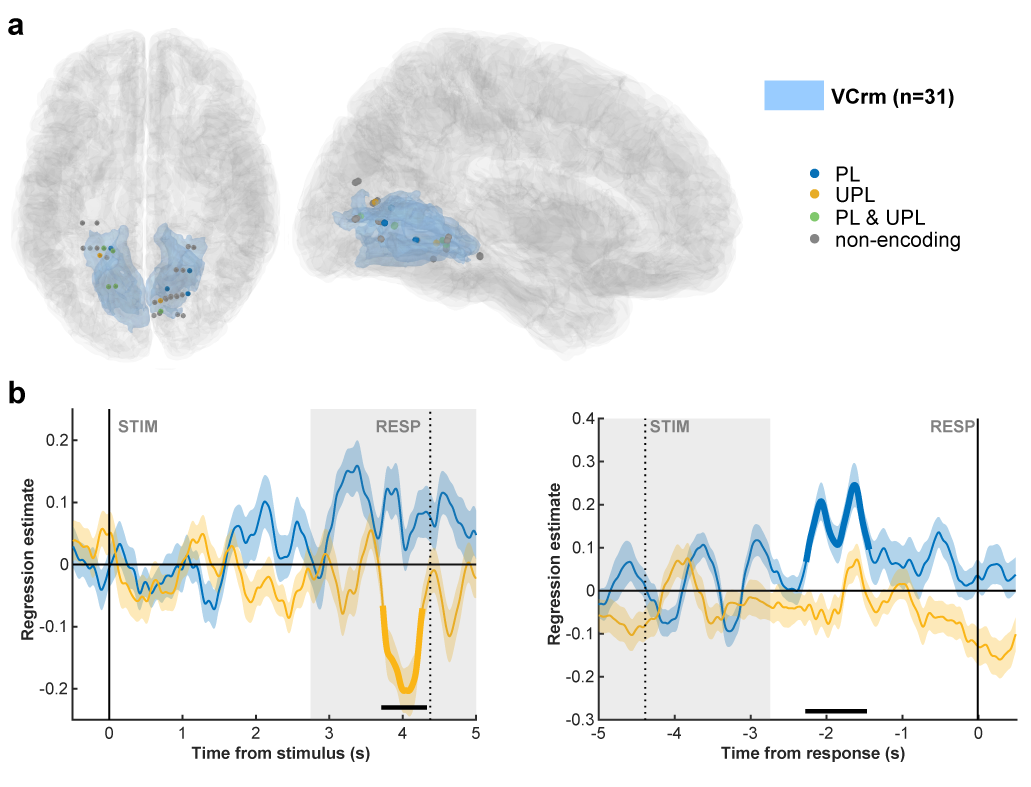

**Figure S3. Delayed and inconsistent relationship between rmVC BGA and unpleasant value.** (a) Anatomical location of all rmVC recording sites projected onto the MNI template brain. Each site is shown as a dot and color-coded by its value-encoding profile (31 sites; 4 PL, 4 UPL, 6 PL&UPL). (b) Group-level results across all sites. We plotted the time-resolved regression estimates of the relationship between trial-wise BGA and item value, averaged across sites and shown separately for pleasant and unpleasant items. Data are locked either to stimulus onset (left panels) or to response onset (right panels). Bold lines indicate significant clusters (p_corr_ < 0.05). Shaded areas denote ± SEM across sites, horizontal bars mark significant clusters (p_corr_ < 0.05). Compared with the aINS (main Fig. 4), this region showed a later-onset cluster in the stimulus-locked analysis, only a few significant UPL sites (n = 4 in panel a), and a transient positive value signal in the response-locked analysis (right panel of b), indicating a weaker and less consistent encoding of unpleasant value.

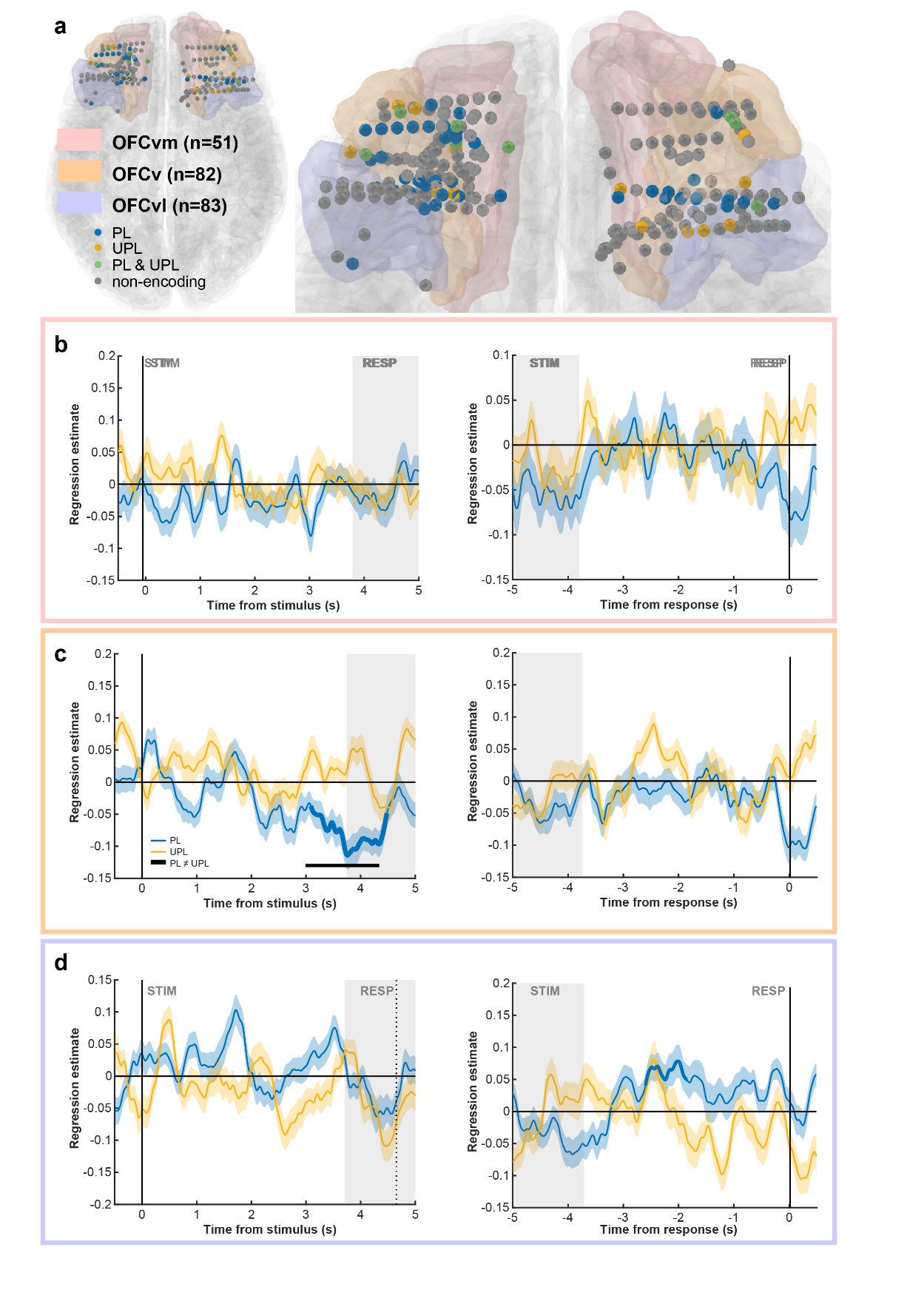

**Figure S4. Orbitofrontal subregions adjacent to the vmPFC do not correlate positively with pleasant value.** (a) Anatomical location of all OFC recording sites projected onto the MNI template brain. Each site is shown as a dot and color-coded by its value-encoding profile. (b-d) Group-level results across all sites. We plotted the time-resolved regression estimates of the relationship between trial-wise BGA and item value, averaged across sites and shown separately for pleasant and unpleasant items. Data are locked either to stimulus onset (left panels) or to response onset (right panels). Bold lines indicate significant clusters (p_corr_ < 0.05). Shaded areas denote ± SEM across sites, horizontal bars mark significant clusters (p_corr_ < 0.05).

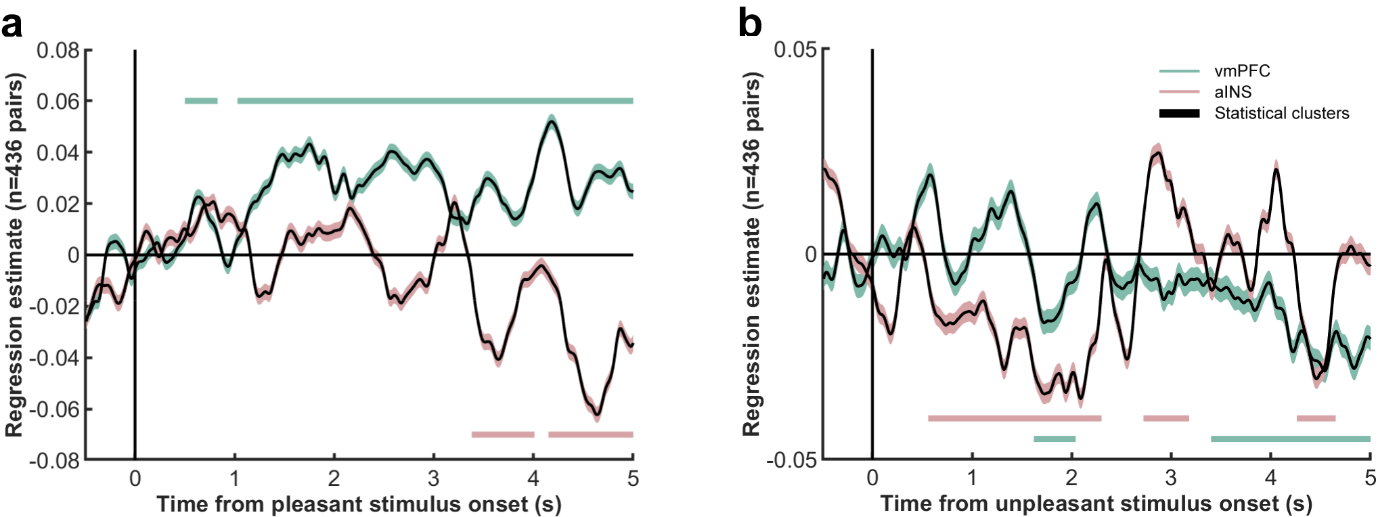

Figure S5. Non-redundant value signals in the vmPFC and aIns during the rating task. Time courses of regression estimates for vmPFC and aIns BGA, included together in a single GLM predicting value and estimated separately for pleasant (a) and unpleasant (b) items. Lines show the mean across all recording sites, and shaded areas indicate ± SEM. Horizontal bars mark significant clusters (p_corr_ < 0.05), highlighting periods when each region contributed non-redundant information about value. By including both regions simultaneously in the model, this analysis demonstrates that the vmPFC and aIns encode value for pleasant and unpleasant items respectively, beyond known effects of their anti-correlation at rest.

### Supplementary tables

| ID | Sex | Age (years) | Hand laterality | Diagnosis age | Suspected epileptic focus |
| --- | --- | --- | --- | --- | --- |
| G1 | M | 38 | R | 35 | Left medial temporal/amygdala/hippocampus |
| G2 | F | 25 | L | 13 | Right insulo-opercular |
| G3 | M | 21 | L | 16 | Undefined |
| G4 | F | 25 | L | 13 | Right insulo-opercular |
| G5 | M | 25 | R | 16 | Left antero-medial temporal |
| G6 | M | 32 | R | 20 | Right parieto-occipital |
| G7 | F | 38 | R | 30 | Left Broca's area (frontal opercular) |
| G8 | M | 56 | R | 47 | Left antero-medial temporal |
| G9 | F | 52 | R | 38 | Right temporo-insulo-opercular |
| G10 | M | 28 | R | 22 | Right parieto-central |
| G11 | M | 39 | R | 35 | Bilateral temporo-insular |
| G12 | M | 41 | R | 36 | Bilateral temporal |
| G13 | F | 18 | L | 16 | Left frontal mesial premotor |
| L1 | F | 32 | R | 11 | NA |
| P1 | F | 37 | L | 24 | Right temporal - pole and superior temporal gyrus |
| P2 | F | 38 | R | 13 | Bilateral frontal |
| P3 | F | 46 | R | 2 | Right frontal medial |
| R1 | M | 29 | L | 5 | NA |
| R2 | F | 35 | R | 14 | NA |
| R3 | F | 26 | R | 12 | NA |
| T1 | F | 30 | R | ? | Right temporo-mesial |
| T2 | F | 22 | R | 17 | Left olfactory gyrus |
| T3 | F | 21 | R | 9 | NA |
| T4 | F | 37 | R | ? | Left hippocampus |
| T5 | M | 43 | R | 37 | Right antero-lateral temporal |
| T6 | F | 41 | L | 34 | Right medial temporal |
| T7 | M | 19 | R | 10 | NA |

Table S1. Demographic and clinical details.

M: male; F: female; L: left; R: right. Diagnosis age refers to the age at which the patient was diagnosed with epilepsy.

| Chosen | | | | | Unchosen | | |
| --- | --- | --- | --- | --- | --- | --- | --- |
| ROI | T-statistic | P-value |  |  | ROI | T-statistic | P-value |
| SPCm | 3,40 | 0,0018 |  |  | pINS | -2,68 | 0,0082 |
| SPC | 2,89 | 0,0055 |  |  | PCC | -3,12 | 0,0036 |
| PFCvm | 2,68 | 0,0090 |  |  | MTCr | -3,38 | 0,0009 |
| pINS | 2,20 | 0,0291 |  |  | IPCv | -3,43 | 0,0009 |
|  |  |  |  |  | PFcdl | -2,75 | 0,0091 |
| Pfrdli | -3,35 | 0,0013 |  |  |  |  |  |
| OFCv | -3,26 | 0,0016 |  |  |  |  |  |
| Hippocampus | -3,03 | 0,0032 |  |  |  |  |  |
| OFCvm | -2,34 | 0,0234 |  |  |  |  |  |

Table S2. Summary of all regions responding to chosen and unchosen option values when asked to choose the best of two pleasant stimuli. BGA in the [-3 0 s] before response was regressed against value across trials for each site within each ROI and significance was tested across beta estimates across all contacts of each ROI.

| Chosen | | | | | Unchosen | | |
| --- | --- | --- | --- | --- | --- | --- | --- |
| ROI | T-statistic | P-value |  |  | ROI | T-statistic | P-value |
| PFCvm | 3,32 | 0,0014 |  |  | PFCvm | 5,29 | 0,0000 |
| Hippocampus | 2,81 | 0,0060 |  |  | MTCc | 5,23 | 0,0000 |
| OFCvl | 2,58 | 0,0115 |  |  | PFcdl | 4,54 | 0,0001 |
| Pfrdls | 2,56 | 0,0128 |  |  | PMrv | 4,34 | 0,0000 |
| MTCr | 2,03 | 0,0440 |  |  | MTCr | 3,91 | 0,0001 |
|  |  |  |  |  | PCC | 3,80 | 0,0005 |
| VCl | -2,72 | 0,0076 |  |  | PMdl | 2,61 | 0,0127 |
|  |  |  |  |  | ITCr | 2,52 | 0,0135 |
|  |  |  |  |  | OFCv | 2,32 | 0,0229 |

Table S3. Summary of all regions responding to chosen and unchosen option values when asked to choose the worst of two pleasant stimuli. BGA in the [-3 0 s] before response was regressed against value across trials for each site within each ROI and significance was tested across beta estimates across all contacts of each ROI.

| Chosen | | | | | Unchosen | | |
| --- | --- | --- | --- | --- | --- | --- | --- |
| ROI | T-statistic | P-value |  |  | ROI | T-statistic | P-value |
| Sv | 5,20 | 0,0000 |  |  | aINS | -4.35 | 2.10^-05^ |
| PMdl | 3,15 | 0,0031 |  |  | Sv | -3,14 | 0,0025 |
| ITCr | 2,78 | 0,0066 |  |  | STCc | -3,11 | 0,0027 |
| VCl | 2,73 | 0,0074 |  |  | pINS | -2,80 | 0,0057 |
| aINS | 2,71 | 0,0075 |  |  | OFCvl | -2,68 | 0,0088 |
| MTCc | 2,45 | 0,0155 |  |  | PFrvl | -2,66 | 0,0096 |
| PFrvl | 2,01 | 0,0484 |  |  |  |  |  |
| Hippocampus | -3,60 | 0,0005 |  |  |  |  |  |
| PFCvm | -2,50 | 0,0146 |  |  |  |  |  |
| Pfrdls | -2,14 | 0,0363 |  |  |  |  |  |

Table S4. Summary of all regions responding to chosen and unchosen option values when asked to choose the best of two unpleasant stimuli. BGA in the [-3 0 s] before response was regressed against value across trials for each site within each ROI and significance was tested across beta estimates across all contacts of each ROI.

| Chosen | | | | | Unchosen | | |
| --- | --- | --- | --- | --- | --- | --- | --- |
| ROI | T-statistic | P-value |  |  | ROI | T-statistic | P-value |
| pINS | -4,57 | 0,0000 |  |  | VCcm | 3,74 | 0,0006 |
| aINS | -3,69 | 0,00029 |  |  | IPCv | 3,32 | 0,0013 |
| VCcm | -3,55 | 0,0011 |  |  |  |  |  |
| STCc | -3,11 | 0,0027 |  |  | Hippocampus | -5,20 | 0,0000 |
| Sv | -2,94 | 0,0045 |  |  | PMrv | -5,19 | 0,0000 |
| ITCm | -2,40 | 0,0194 |  |  | MTCr | -3,97 | 0,0001 |
| ITCr | -2,10 | 0,0389 |  |  | Mdl | -3,48 | 0,0009 |

Table S5. Summary of all regions responding to chosen and unchosen option values when asked to choose the worst of two unpleasant stimuli. BGA in the [-3 0 s] before response was regressed against value across trials for each site within each ROI and significance was tested across beta estimates across all contacts of each ROI.
